# Low-intensity focused ultrasound to the human insular cortex differentially modulates the heartbeat-evoked potential: a proof-of-concept study

**DOI:** 10.1101/2024.03.08.584152

**Authors:** Andrew Strohman, Gabriel Isaac, Brighton Payne, Charles Verdonk, Sahib S. Khalsa, Wynn Legon

## Abstract

**Background:** The heartbeat evoked potential (HEP) is a brain response time-locked to the heartbeat and a potential marker of interoceptive processing. The insula and dorsal anterior cingulate cortex (dACC) are brain regions that may be involved in generating the HEP. Low-intensity focused ultrasound (LIFU) is a non-invasive neuromodulation technique that can selectively target sub-regions of the insula and dACC to better understand their contributions to the HEP.

**Objective:** Proof-of-concept study to determine whether LIFU modulation of the anterior insula (AI), posterior insula (PI), and dACC influences the HEP.

**Methods:** In a within-subject, repeated-measures design, healthy human participants (n=16) received 10 minutes of stereotaxically targeted LIFU to the AI, PI, dACC or Sham at rest during continuous electroencephalography (EEG) and electrocardiography (ECG) recording on separate days. Primary outcome was change in HEP amplitudes. Relationships between LIFU pressure and HEP changes were examined using linear mixed modelling. Peripheral indices of visceromotor output including heart rate and heart rate variability (HRV) were explored between conditions.

**Results:** Relative to sham, LIFU to the PI, but not AI or dACC, decreased HEP amplitudes; this was partially explained by increased LIFU pressure. LIFU did not affect time or frequency dependent measures of HRV.

**Conclusions:** These results demonstrate the ability to modulate HEP amplitudes via non-invasive targeting of key interoceptive brain regions. Our findings have implications for the causal role of these areas in bottom-up heart-brain communication that could guide future work investigating the HEP as a marker of interoceptive processing in healthy and clinical populations.

## INTRODUCTION

Interoception refers to an internal sense of the physiologic conditions of the body and relies on the transmission of neural signals between the brain and internal organs like the heart, lungs, gut, and vasculature [1–4]. Dysfunction in interoceptive processing has been broadly implicated in mental health and chronic pain disorders [3–5], making efforts to modulate interoceptive signaling potentially promising for future therapeutic interventions.

Afferent information from internal organs travel via the vagus and glossopharyngeal nerves to the brainstem and thalamic targets and integrate with lamina I spinothalamic projections before relaying to the insula and dorsal anterior cingulate cortex (dACC) [1,2,6–9]. Both the PI and the dACC receive direct afferent interoceptive information through two distinct thalamic nuclei while the AI communicates with PI and dACC without directly receiving this information, suggesting complementary but specific roles in interoceptive processing [1,7,10–12]. This makes the insular subregions and the dACC intriguing targets for both the pathophysiologic investigation and neuromodulation therapies across a range of mental health and chronic pain conditions that may involve brain-body dysregulation [13–19].

The heartbeat-evoked potential (HEP) is a brain-derived electrophysiologic signal time-locked to each heartbeat, that may indicate the central representation of signaling from the heart and baroreceptors [20–23]. Alterations in the HEP have been implicated in chronic pain [24] and mental health disorders like anxiety and depression [21,25–27], making the HEP a potential therapeutic biomarker across a wide range of clinical conditions. Intracranial recordings have also established the anterior insula (AI), posterior insula (PI), and dACC as either generating or contributing to the HEP [20,28–30], but it is presently unknown whether non-invasive focal neuromodulation can impact the HEP.

Low-intensity focused ultrasound (LIFU) is an increasingly investigated non-invasive neuromodulatory approach that leverages mechanical energy to transiently alter brain activity at a millimeter-sized focus with an adjustable depth [31,32]. LIFU has been demonstrated to affect electrophysiologic, bold oxygen-level dependent (BOLD), and neurotransmitter activity for both superficial and deep structures in humans that translates to behavior changes [33–39]. More specifically, recent work from our lab demonstrates LIFU can specifically target the PI, AI, and dACC with differential effects on subjective report of pain, pain-evoked electrophysiologic potentials, and heart rate variability [40–42].

The purpose of this proof-of-concept study was to investigate site-specific roles of the PI, AI and dACC on the HEP, heart-rate and heart-rate variability (HRV), using LIFU to focally target these regions. We conducted this study during physiological resting conditions as many clinical brain stimulation therapies are applied at rest and evidence suggests dysfunction in neural and cardiac activity at rest across numerous clinical disorders [24,43–45]. We hypothesized LIFU would confer an inhibitory effect based on prior findings [40,41], and specifically LIFU to the PI and dACC to attenuate the HEP amplitude as both regions receive cardiac afferents via the thalamus [1,7,12]. By contrast, the AI does not receive direct cardiac input [1,14], thus we hypothesized that LIFU to the AI would not affect the HEP amplitude as compared to sham.

## MATERIALS AND METHODS

### Participants

The Institutional Review Board at Virginia Tech approved all experimental procedures (IRB #21-796). N=16 healthy participants (25.7 years ± 3.4 years; range (20-34); M/F 5/11), who met all inclusion/exclusion criteria provided written informed consent to all aspects of the study. Inclusion criteria were males and females ages 18-65 while exclusion criteria included contraindications to imaging (magnetic resonance imaging (MRI) and computed tomography (CT)), a history of neurologic disorder or head injury resulting in loss of consciousness for >10 minutes or drug dependence, and any active medical disorder or current treatment with potential central nervous system effects and no history of chronic pain.

### Overall Study Design and Timeline

All participants completed a total of five visits in this sham-controlled cross-over design study. Following informed consent, the first visit comprised a structural brain MRI and CT for the purpose of LIFU targeting and acoustic modelling (see below). The remainder of the visits were formal testing sessions of LIFU to the anterior insula, posterior insula, dorsal anterior cingulate or Sham, randomized within and between subjects. A minimum of two days was required between visits to mitigate any potential carry-over effects [46].

This data was extracted from a larger study investigating LIFU modulation of pain processing [42]. For the present study, we investigated the HEP during 10 minutes of LIFU application when participants were not completing any task. **Figure 1** shows the LIFU conditions and LIFU delivery during the 10 minutes.

**Figure 1.**
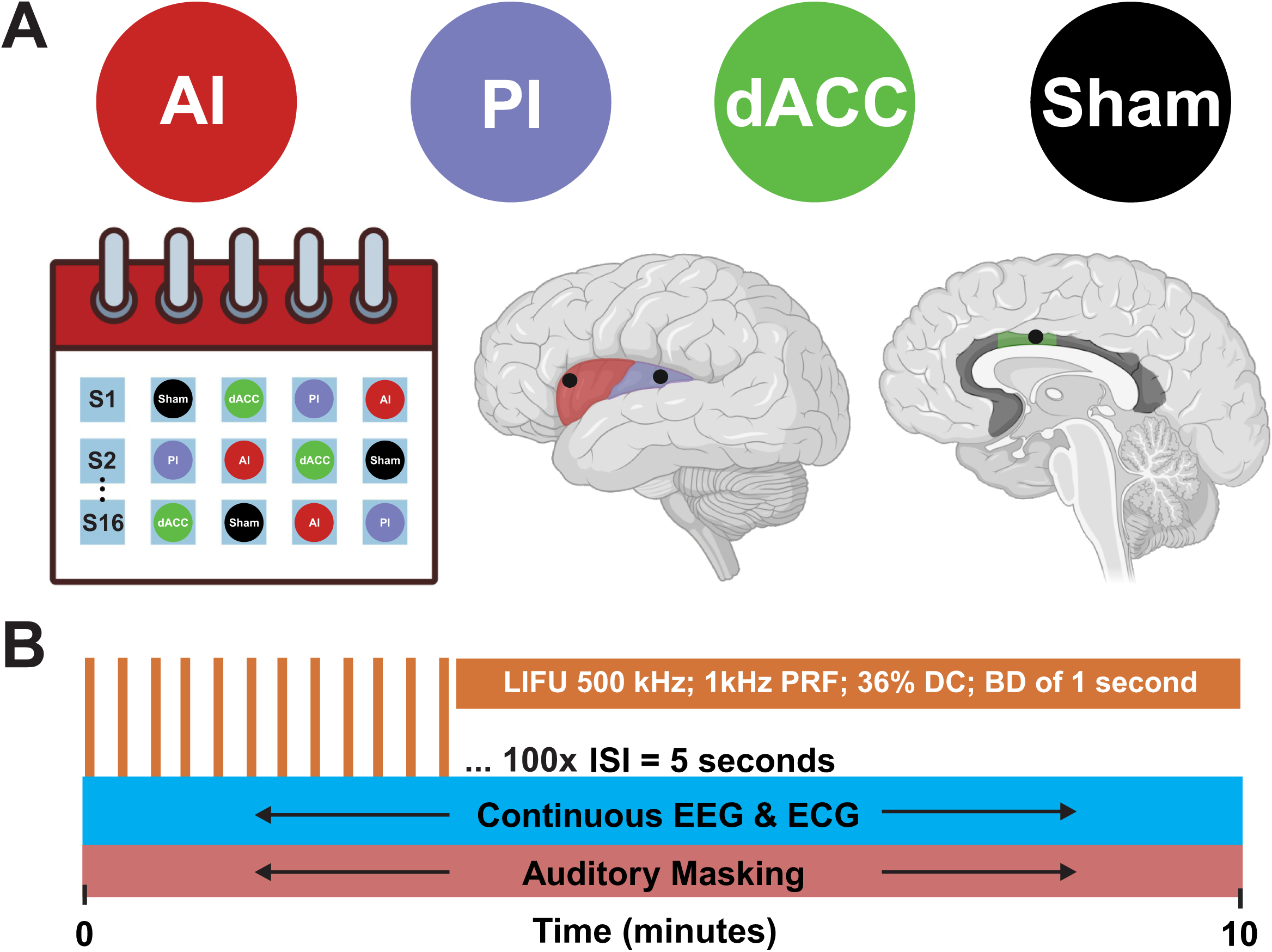
Study Design. **A.** Graphical depiction of formal testing days. Participants (N = 16) completed four arms randomly assigned on four separate days: Low-intensity focused ultrasound (LIFU) to the anterior insula (AI), posterior insula (PI), dorsal anterior cingulate cortex (dACC) or Sham. Black dots on brains represent approximate locations of each target for the AI (left dorsal anterior short gyrus), PI (left dorsal posterior long gyrus), and dACC (Montreal Neurologic Institute (MNI) coordinates [0,18,30]) for all participants. Graphical brains were adapted from Biorender.com. **B.** For each session, data was collected over a 10 minute time window while participants sat at rest during continuous electroencephalogram (EEG) and electrocardiogram (ECG) recording. For each active day, a total of 100 LIFU stimulations were delivered over the 10-minute time window with a fixed interstimulus interval (ISI) of 5 seconds. Continuous auditory masking was delivered over the entire time window for all conditions. Blow-up of a single orange bar details the LIFU stimulation parameters. One second of LIFU was delivered with a fundamental frequency of 500 kHz, a pulse repetition frequency (PRF) of 1 kHz, a duty cycle (DC) of 36%, and a total burst duration (BD) of 1 second.

### MRI and CT Acquisition

MRI data were acquired on a Siemens 3T Prisma scanner (Siemens Medical Solutions, Erlangen, Germany) at the Fralin Biomedical Research Institute’s Human Neuroimaging Laboratory. Anatomical scans were acquired using a T1-weighted MPRAGE sequence with a TR = 1400 ms, TI = 600 ms, TE = 2.66 ms, flip angle = 12 (degrees), voxel size = 0.5x0.5x1.0 mm, FoV read = 245 mm, FoV phase of 87.5%, 192 slices, ascending acquisition. CT scans were collected with a Kernel = Hr60 in the bone window, FoV = 250 mm, kilovolts (kV) = 120, rotation time = 1 second, delay = 2 seconds, pitch = 0.55, caudocranial image acquisition order, 1.0 mm image increments for a total of 121 images and scan time of 13.14 seconds.

### Data Acquisition

*Electroencephalography (EEG)*. Surface EEG was collected using a DC amplifier (GES 400, Magstim EGI, Eugene, OR, USA) and Net Station ^TM^ 5.4 EEG software at a sampling rate of 1 kilohertz (kHz) with a 10 mm silver-silver chloride cup electrode placed on the vertex (Cz), referenced to the right mastoid. In this proof-of-concept study, we measured EEG signals using a single electrode placed at the Cz location. The Cz site was selected on the basis that 1) this location is one of the most commonly identified in meta-analytic studies of the HEP [21], and 2) Cz is a common site chosen in single electrode studies of visceral-evoked potentials [47–49]. The scalp was first prepped with a mild abrasive gel (NuPrep; Weaver and Company, Aurora, CO) and rubbing alcohol. The electrode was filled with a conductive paste (Ten20 Conductive; Weaver and Company, Auora, CO) and secured with medical tape. Electrode impedances were verified (<50 kΩ) before recording data, which was stored on a PC for offline analysis.

*Electrocardiography (ECG).* Two ECG electrodes (MedGel^TM^, MDSM611903) were placed on the anterior surface of both forearms immediately distal to the antecubital fossa (Einthoven lead I configuration) sampled at 1 kHz using a physiologic data acquisition box (Physio16, Magstim EGI, OR, USA) that has an integrated ground (GES 400, Magstim EGI, Eguene, OR, USA). Data was stored on a PC for offline analysis.

### LIFU Transducers and Waveform

Two different transducers were used based upon individual participant head morphology and depth of target. For insula targets, a single-element 500 kHz transducer (Sonic Concepts H-281) was used with an active diameter of 45.0 mm and 38.0 mm focal length from the exit plane. The transducer also had a solid water coupling over the radiating surface to the exit plane. To target the dACC, a single-element 500 kHz transducer (Sonic Concepts H-104) was used with an active diameter of 64.0 mm and 52.0 mm focal length from the exit plane.

Focused ultrasound waveforms were generated using a dual channel function generator (BK 4078B Precision Instruments). Channel 1 was a 5Vp-p square wave burst of 1 kHz (N=1000) with a duty cycle of 36% used to gate channel 2 that was a 500 kHz sine wave. Channel 2 output was sent through a 100-W linear RF amplifier (E&I 2100L; Electronics & Innovation) before being sent to the transducers. LIFU was applied for 100 stimulations (burst duration = 1 sec) separated by a fixed inter-stimulus interval (ISI) of 5 seconds for a total application time of 10 minutes. For both AI and PI, the applied external pressure was 380 kPa with a spatial peak-pulse average intensity (I_sppa_) of 4.2 W/cm^2^, a spatial peak temporal average (I_spta_) of 1.5 W/cm^2^ and a mechanical index of (MI) of 0.2. For dACC, the applied external pressure was 400 kPa with an I_sppa_ of 4.5 W/cm^2^, an I_spta_ of 1.62 W/cm^2^, and an MI of 0.23.

### LIFU Targeting and Application

The transducer was coupled to the head using conventional ultrasound gel and our custom mineral oil/polymer coupling pucks [50]. These pucks have negligible attenuation at 500 kHz [50] and can be made with varying stand-off heights that allow for precise axial (depth) targeting based on individual insular and dACC target depths (**Table A.1**). For both AI, PI, and dACC conditions, an appropriate coupling puck was made so that the focal spot of the transducer was exactly overlaid on the insular or dACC target. Each participant’s left AI (dorsal anterior short gyrus) and PI (dorsal posterior long gyrus) target was identified with the aid of an insular atlas [46] and depth was measured from the scalp. For dACC, MNI coordinate [0,18,30] was used based upon Neurosynth reverse inference software for the term ‘pain’ and prior literature on the fMRI signature of heat pain [51,52]. Placement of the transducer on the scalp for each target site was aided using a neuronavigation system (BrainSight, Rogue Research, Montreal, QUE, CAN). LIFU was only delivered if placement error on the scalp was < 3 mm.

### Auditory Masking

Participants were given headphones connected to a tablet that had a multi-tone white noise generator app. There were a variety of sound options that could be mixed and layered on top of each other to create a multitone that has been established to effectively mask a 1 kHz pulsed ultrasound soundwave [53]. The volume was set to a level where participants could not hear normal conversations with the intensities ranging from 70 to 75 dB.

### Questionnaires

#### Review of Symptoms

A review of symptoms questionnaire [54] asked about the presence of various symptoms and their severity (absent / mild / moderate / severe) scored on a scale of 0-3. This questionnaire was collected at the beginning of each visit and 30 minutes after the LIFU or Sham application in order to indicate any changes of symptoms from the intervention. Counts for each symptom at each severity level were calculated for both pre and post LIFU for each condition (AI, PI, dACC, Sham). Counts are also calculated adjusting for the presence of symptoms after LIFU as compared to symptoms before LIFU to better understand if new symptoms occurred after the intervention to each target site.

#### Auditory Masking

An auditory masking questionnaire [50,55] was administered at the end of each visit (sessions 2 – 5) with the following questions: “I could hear the LIFU stimulation”, “I could feel the LIFU stimulation”, and “I believe I experienced LIFU stimulation.” Participants were asked to rate each question on a 7-point Likert scale: strongly disagree / disagree / somewhat disagree / neutral / somewhat agree / agree / strongly agree. To statistically test for differences between conditions, scores for each question (“Hear”, “Feel”, “Believe”) were converted to numerical values from 0-6 and compared across conditions using a kruskal-wallis test at a significance level of p < 0.05. Data are reported as the mean ± SEM for each condition.

### Formal Testing Days

On each formal testing day, participants were seated in a comfortable chair and connected to continuous EEG and ECG (see above). After the initial set of tasks prior to LIFU (see In et al., 2024 [42]), LIFU was stereotaxically targeted to either the AI, PI, dACC or Sham. For Sham, the transducer was randomly positioned at one of those sites within and between participants. Participants were also given headphones to play the masking auditory stimulus throughout the entire LIFU application period (**Figure 1A**). As outlined above, LIFU was delivered with a burst duration of 1 second and an ISI of 5 seconds for a total of 100 stimulations for a total duration of 10 minutes (**Figure 1A&B**). The beginning and end of LIFU stimulation denoted the EEG and ECG data of interest for pre-processing and analysis.

### Data Pre-Processing

#### Electrocardiography

ECG data was first filtered from 0.3-35 Hz using a 3^rd^ order Butterworth filter and movement artifacts were manually removed. R-R peaks were then extracted using the automated detection program *findpeaks* in Matlab and confirmed manually. Only R-R peaks for the time window during LIFU application were used for data analysis.

#### Electroencephalography

EEG pre-processing was adapted from Petzschner et al. (2019) [56]. EEG data were preprocessed using custom scripts written in Matlab 2022a® (The MathWorks, Inc., Natick, MA). Data were band-pass filtered from 0.3-35 Hz using a third-order Butterworth filter and the *filtfilt* function. The 0.3 Hz lower bound for the bandpass filter was chosen based on previous literature suggesting higher high-pass values can introduce artifact into the signal [56–58]. EEG data were epoched around each R-peak from 100 ms before to 667ms after the R-peak. 667 ms was chosen as this was the maximum value that still included individuals with a resting heart-rate < 90 bpm. Trials were rejected for either of two reasons: 1) If greater than two percent of the data across the epoch’s time window exceeded three standard deviations of the mean HEP amplitude or 2) If the instantaneous heart rate at any time exceeded 90 beats per minute (bpm) for that trial. In order to avoid artifact from preceding heartbeats, we did not baseline correct our epochs as previously performed [56]. Based on the above criteria, the median (interquartile range (IQR)) number of trials removed was 52.5 (35), 56.5 (18), 57 (49.5), and 54 (22) for AI, PI, dACC, and Sham conditions, respectively (range: 7-416). A Kruskal-Wallis test revealed no significant difference between conditions (Chi-square = 0.27(3), p = 0.98). The total number of trials removed accounted for 12.16% of the data across all participants and conditions. This is a similar percentage of removed trials compared to prior literature [21].

The mean ± SD final number of trials included in the analysis was 539.6 ± 85.9, 500.5 ± 69.1, 510.8 ± 70.1, and 529.9 ± 73.1 for AI, PI, dACC, and Sham conditions, respectively. A one-way ANOVA revealed no significant difference between conditions (F = 1.70 (3,45), p-value = 0.18). The time window of interest (TOI) for HEP quantification was 200 to 667ms after the R-peak. This was based on Coll et al. (2021) as 200ms post R-peak is the earliest time HEP effects have been reported, and which corresponds to the time of increased baroreceptor firing when the systolic blood outflow stretches the walls of the aortic arch and carotid sinus [21,59]. In addition, the influence of cardiac field artifact (CFA) on HEP amplitude is presumed to be limited from 200ms post R-peak [21,56,60]. The final time point follows the approximate last time point where HEP effects have been previously reported and is based on a maximum resting heart rate of 90 bpm [56].

### Data Analysis

#### Acoustic Modelling

Computational models were developed using individual subject MR and CT images to evaluate the wave propagation of LIFU across the skull and the resultant intracranial acoustic pressure maps. Simulations were performed using the k-Wave MATLAB toolbox [61], which uses a pseudospectral time domain method to solve discretized wave equations on a spatial grid. CT images were used to construct the acoustic model of the skull, while MR images were used to target LIFU at either the AI, PI, or dACC target, based on individual brain anatomy. Details of the modeling parameters can be found in Legon et al. (2018) [62]. CT and MR images were first co-registered and then up-sampled for acoustic simulations and the acoustic parameters for simulation were calculated from the CT images. The skull was extracted manually using a threshold intensity value and the intracranial space was assumed to be homogenous as ultrasound reflections between soft tissues are small [63]. Acoustic parameters were calculated from CT data assuming a linear relationship between skull porosity and the acoustic parameters [64,65]. The computational model of the ultrasound transducer used in simulations was constructed to recreate empirical acoustic pressure maps of focused ultrasound transmitted in the acoustic test tank similar to previous work [55,62,66,67]. For each subject and each stimulation site, the estimated LIFU pressure at the beam focus in kilopascals (kPa) was extracted and used for later analysis.

#### HEP Time Domain Analysis

To characterize LIFU-induced changes in HEP amplitude, we utilized non-parametric permutation testing with custom Matlab™ scripts following procedures recommended by Maris and Oostenveld (2007) [68]. We used the F-statistic from a one-way ANOVA with the four conditions (AI,PI, dACC, Sham) across the epoched EEG time window (100ms before to 667ms post R-peak), with 1000 permutations at each millisecond at a significance level of p< 0.05. Data is displayed without any cluster threshold in order to visualize any significant time points. The effect size for the average HEP amplitude across each significant time window is reported as the eta-squared (^2^). Time points before 200ms post R-peak (despite being in the window of the CFA) were included in the analysis to identify any time points with significant differences within the CFA. Any significant findings were used for control analyses (see below) to test whether these differences explain observed differences in the HEP (200 – 667 ms). Post-hoc comparisons on the average amplitude for each time window with significant differences (p < 0.05) in the TOI of the HEP were explored using Tukey’s test.

#### Control Analyses

Control analyses were adopted from Petzschner et al. (2019), with a couple of additions [56]. We wanted to rule out that changes in HEP amplitude were not caused by changes in cardiac electrical activity, as measured by the ECG, which may propagate throughout the body and affect EEG scalp potentials. For this purpose, we epoched the ECG data using the same approach as for the EEG data (see above), and we tested the condition effect (AI vs. PI vs. dACC vs. Sham) on the averaged ECG waveform. To this end, the F-statistic from a one-way ANOVA with the four conditions (AI, PI, dACC, Sham) was tested across the entire window (- 100ms to +667ms around the R-peak) using 1000 permutations at each millisecond at a significance level of p < 0.05. Data is displayed without any cluster threshold in order to visualize any significant time point differences in the ECG.

As an additional check to rule out potential confounding cardiac effects, we extracted ECG amplitudes from the same time window as any observed significant findings in the HEP data. We then employed linear regression on the mean value from the time windows, testing whether the change in amplitude in the HEP data was explained by any change in amplitude in the ECG data at these specific time points. Data is reported as the adjusted R^2^, F-statistic, and p-value at a significance level of p < 0.05.

For any significant findings in the EEG data during the CFA, the same approach was used as above with the ECG data. We averaged across all significant time points within the CFA time window, subtracted across the conditions identified from post-hoc testing in the EEG data, and employed linear regression to test whether the change in amplitude in the HEP window was explained by any change in amplitude in the CFA window. Data is reported as the adjusted R^2^, F-statistic, and p-value at a significance level of p < 0.05.

#### Analysis of Pressure

Estimated LIFU pressures for each subject at each target were tested across conditions to evaluate for significant differences using a one-way ANOVA with factor CONDITION (AI, PI, and dACC) at a significance level of 0.05. With the presence of significant differences, we employed linear mixed effect modelling to help explain any observed differences in the HEP time windows identified during permutation testing. We ran four models using the differences in the HEP amplitude taken between conditions based on post-hoc testing from permutation analysis as the dependent variable. We then included either pressure alone, condition alone, pressure and condition, or pressure, condition, and the pressure-by-condition interaction as fixed effects for each of the four models, respectively. Subjects were used as a random effect in all models. The theoretical likelihood ratio test was used to determine the best model and is reported with the log likelihood for each model along with the LRstat (deltaDF) and the p-value at a significance level of 0.05. The model with the lowest Akaike Inference Criterion (AIC) was used if the theoretical likelihood ratio test was not significant. The best overall model is reported with the F-statistic (DF), the p-value at a significance level of 0.05, and the adjusted R^2^ (R^2^_adj_). The beta coefficients (95% CI), t-statistic (DF), and p-values for each variable are also reported.

#### Analysis of Heart Rate & Heart Rate Variability

We further explored whether LIFU had any effects on heart rate and heart rate variability (HRV), defined as the change in time intervals between consecutive heartbeats [69]. We first calculated median heart rate to investigate if there were any differences between conditions. We used a one-way kruskal-wallis test with main factor CONDITION (AI, PI, dACC, Sham). Data is reported as the median (IQR) with the chi-square (df), p-value at a significance level of p < 0.05, and the eta-squared (^2^). We also calculated four HRV metrics during the period of LIFU application: the standard deviation in normal sinus beats (SDNN), low frequency (LF) power, high frequency (HF) power, and the LF/HF ratio [69]. The SDNN is a time-domain metric and was calculated as the standard deviation of the time in milliseconds between consecutive normal sinus beats. For frequency domain metrics (LF power, HF power, LF/HF ratio), R-R interval data during LIFU or sham application was cubic spline interpolated and transformed using the fast-fourier transform. The LF power (0.04-0.15 Hz) and HF power (0.15-0.4 Hz) bands during LIFU or Sham application were calculated for each subject for each condition [69]. The LF/HF ratio was calculated by dividing the LF by the HF power. All HRV metrics were tested using a one-way ANOVA with main factor CONDITION (AI, PI, dACC, Sham). Data is reported as the mean ± SEM, the F-statistic (df) at a significance level of p < 0.05, and the eta-squared (^2^).

## RESULTS

### Insula & dACC target depths

The mean ± SD depth from the scalp to each target was 34.5 ± 2.6 mm, 39.4 ± 2.6 mm, and 47.3 ± 3.4 mm for the AI, PI, and dACC targets, respectively (**Table A.1**). Individual participant depths and means for each sex can be seen in **Table A.1**.

### Acoustic Modelling

For the insula transducer, the beam profile in free water had a lateral full-width at half maximum (FWHM) resolution in the lateral (XY) resolution ± 1.7 mm at the axial (Z) beam focus and a FWHM axial (YZ) resolution of ± 11.5 mm with a focal depth of 38 mm from the exit plane of the transducer (**Figure 2A**). For the dACC transducer, the beam profile in free water revealed a FWHM lateral (XY) resolution of ± 1.5 mm at the axial (Z) beam focus and a FWHM axial (YZ) resolution of ± 10 mm with a focal depth of 52 mm from the exit plane of the transducer (**Figure 2B**). The volumes these transducers overlaid well with the individual subject target depths (**Table A.1**).

**Figure 2.**
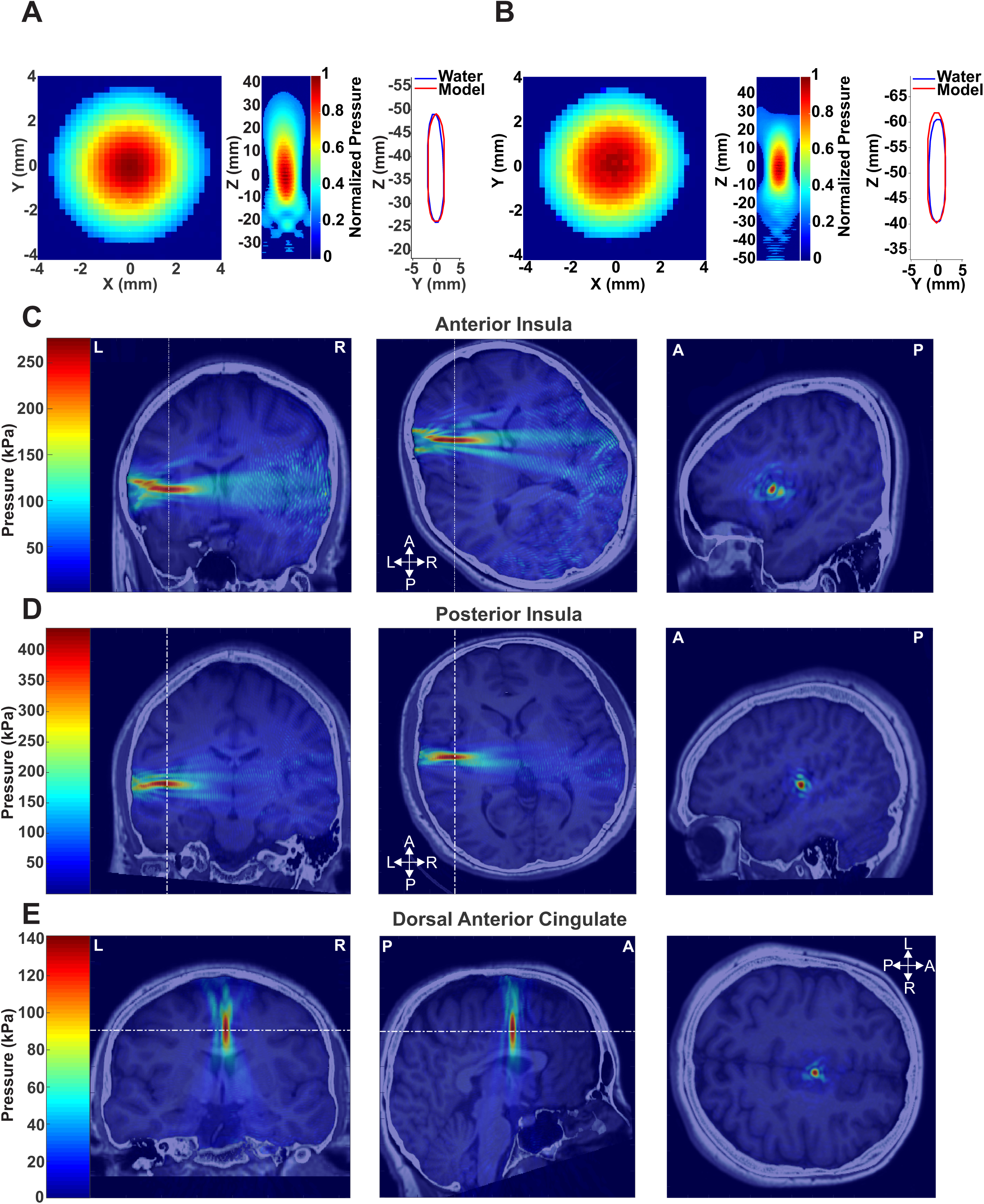
Transducer Characteristics and Acoustic Modelling. **A.** (Left) Pseudocolor XY and YZ empirical measurements showing the normalized pressure map for the transducer used for targeting anterior (AI) and posterior (PI) insula. (Right) Comparison of the YZ full-width half maximum (FWHM) beam profile of the transducer in free water (blue) compared to the YZ beam profile used for acoustic modelling. **B.** (Left) Pseudocolor XY and YZ empirical measurements of the normalized pressure map for the transducer used for targeting the dorsal anterior cingulate cortex (dACC). (Right) Comparison of the YZ beam profile of the transducer in free water (blue) compared to the YZ beam profile of the transducer used for acoustic modelling. **C.** Acoustic models using individual magnetic resonance and computed tomography scans from a representative subject showing 500 kHz LIFU targeting the AI. Color scale is pressure in kilopascals (kPa). Vertical dashed white lines in the coronal and transverse planes represent the slice taken for the sagittal view. **D.** Acoustic models using individual magnetic resonance and computed tomography scans from a representative subject showing 500 kHz LIFU targeting the PI. Color scale is pressure in kPa. Vertical dashed white lines in the coronal and transverse planes represent the slice taken for the sagittal view. **E.** Acoustic models using individual magnetic resonance and computed tomography scans from a representative subject showing 500 kHz LIFU targeting the dACC. Color scale is pressure in kPa. Vertical dashed white lines in the coronal and transverse planes represent the slice taken for the sagittal view.

The axial FWHM comparison between the empirical measurements in free water and the modelled waveform for both transducers demonstrated good agreement validating use in the acoustic models (**Figures 2A&B**). Acoustic models for each site (AI, PI, dACC) from representative subjects can be seen in **Figures 2C-E**.

### HEP Time Domain Analysis

Permutation analysis (permutations = 1000, 1ms intervals, p < 0.05, no cluster threshold) revealed three distinct time windows with significant differences between conditions (**Figure 3A & B**). The first window within the TOI of the HEP (200 – 667 ms) was a 13-millisecond window from 274 to 286 milliseconds after the R-peak. Of the 13ms, 9 non-consecutive time-points were significant and the p-values for the entire 13ms window ranged from 0.038 to 0.057 (^2^ = 0.09) (**Figure 3A & B**). As such, we used the entire window for subsequent post-hoc testing and control analyses. Post-hoc testing revealed significant differences between PI and Sham (p = 0.033) (**Figure 3C**). The second significant window within the TOI of the HEP was a 14ms time window from 648 to 661ms post R-peak (all p-values < 0.05, ^2^ = 0.11) (**Figure 3A & B**). Post-hoc testing revealed a significant difference between PI and dACC conditions (p=0.014), but not Sham (**Figure 3C**). A plot depicting F-statistics and p-values from permutation testing across time can be seen in **Supplemental Figure A.3A**.

**Figure 3.**
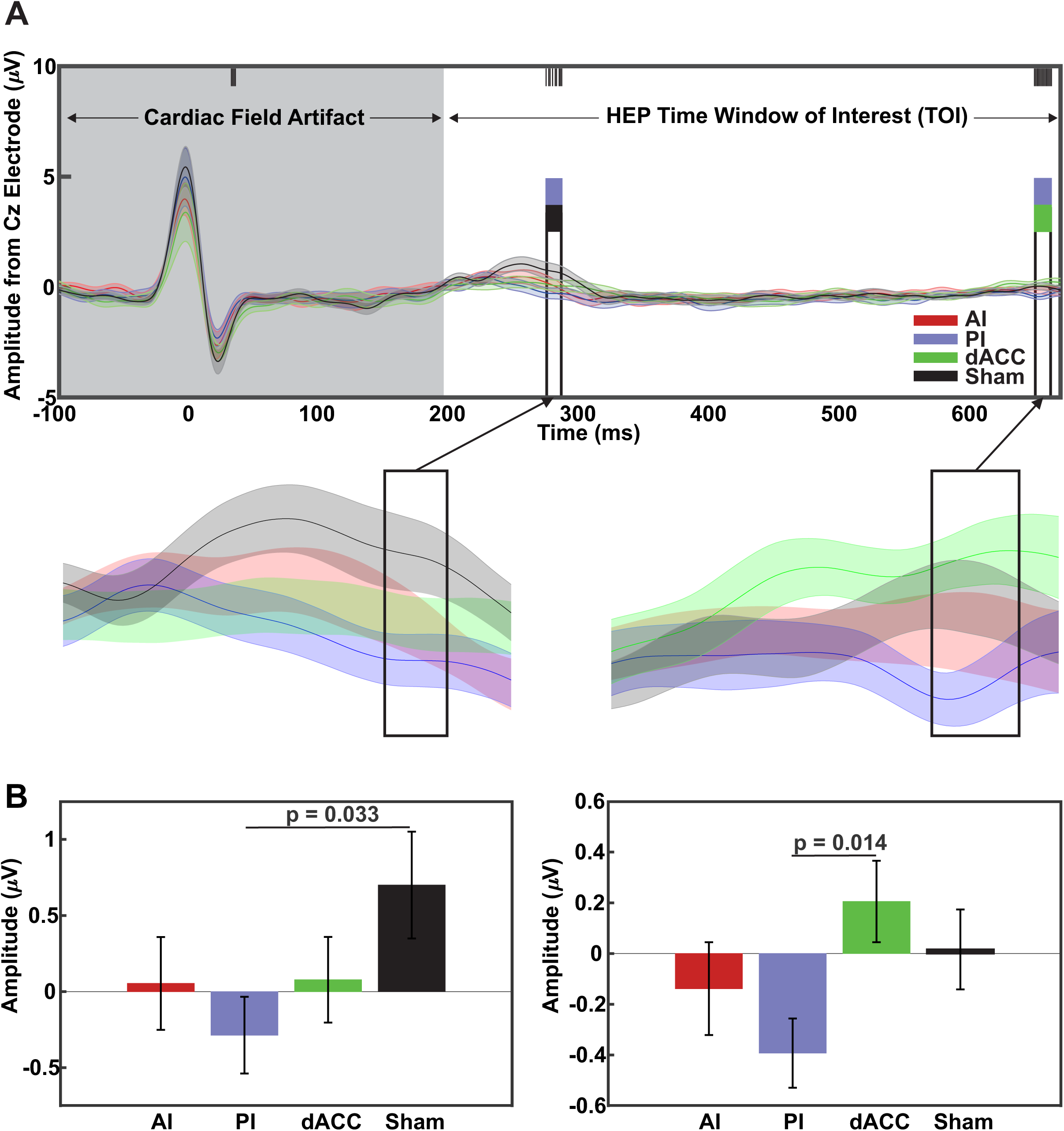
Analysis of HEP amplitudes. **A.** Group (N=16) mean ± SEM heartbeat-evoked potential (HEP) amplitudes across time. X-axis is time in milliseconds (ms) and y-axis is amplitude in microvolts (μV). Grey background represents the period of the cardiac field artifact while the white represents the HEP time window of interest (TOI). Vertical black bars at the top represent significant time points from permutation analysis (1000 permutations, p < 0.05). Vertical colored bars represent significant differences across conditions from post-hoc testing at each HEP time window of interest. F-statistics and p-values from permutation testing can be seen in Supplemental Figure A.3A. (Below) Magnified Group (N=16) mean ± SEM heartbeat-evoked potential (HEP) traces around the time periods of first (left) and second (right) time periods of significant results. Black arrow points to the same significant time window in A. **B.** (Left) Group (N=16) mean ± SEM HEP amplitudes (μV) across conditions within the first significant period in the HEP time window of interest. (Right) Group (N=16) mean ± SEM HEP amplitudes (μV) across conditions within the second significant period in the HEP time window of interest. Horizontal bars and p-values represent results from post-hoc comparisons.

### Control Analyses

In order to test for significant differences in the ECG trace within the TOI of the HEP, we ran the same permutation analysis (permutations = 1000, 1ms intervals, p < 0.05, no cluster threshold) across groups in the ECG data. There were no significant differences across any of the conditions (**Figure 4A**). A plot depicting F-statistics and p-values from permutation testing across time can be seen in **Supplemental Figure A.3B**.

**Figure 4.**
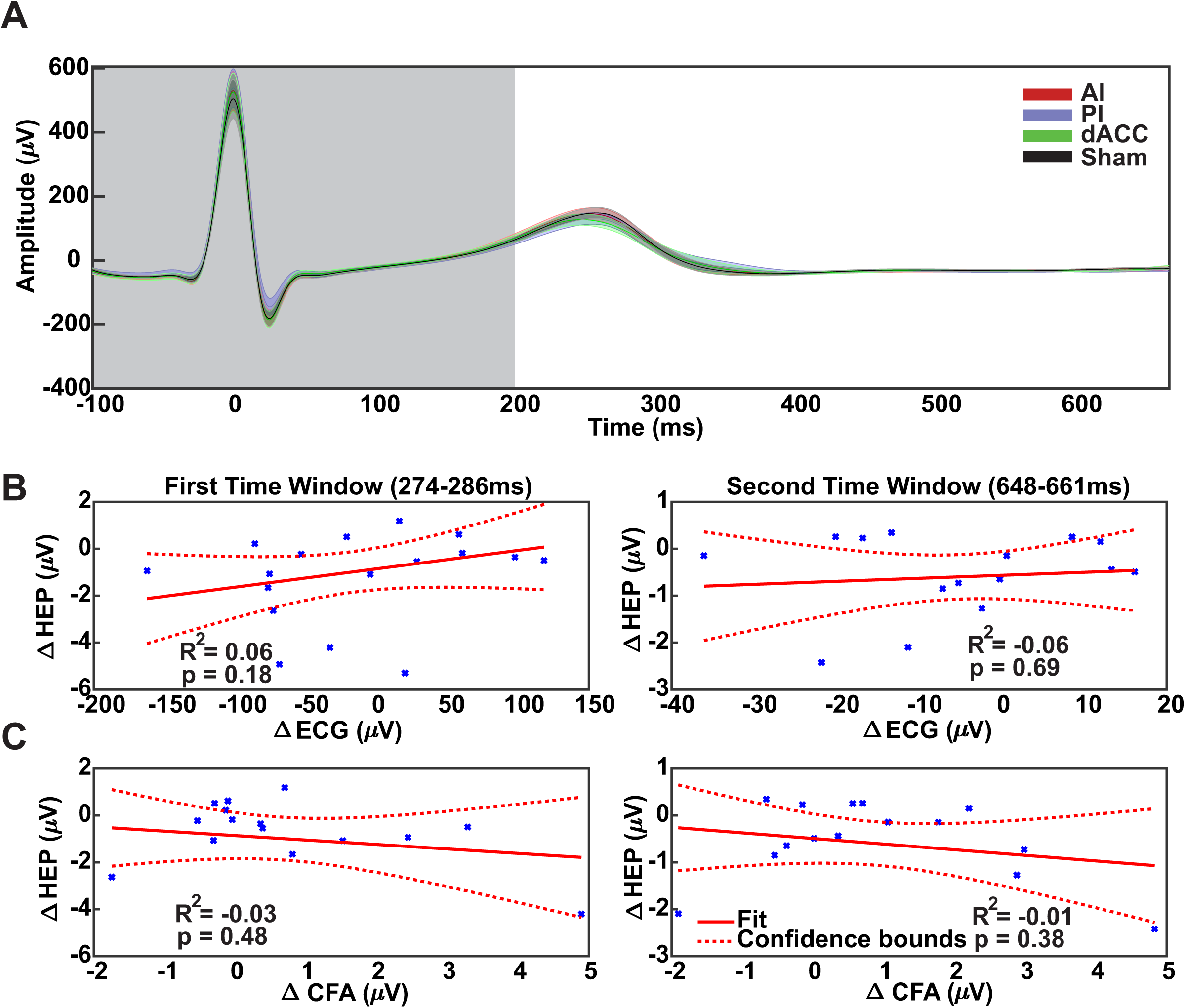
Control analyses on ECG and cardiac field artifact. **A.** Group (N=16) mean ± SEM electrocardiogram (ECG) amplitudes across time. X-axis is time in milliseconds (ms) and y-axis is amplitude in microvolts (μV). Grey background represents the equivalent period of the cardiac field artifact from EEG data while the white represents the equivalent heartbeat-evoked potential (HEP) time window of interest. F-statistics and p-values from permutation testing can be seen in Supplemental Figure A.3B. **B.** (Left) Linear regression comparing the average change in the HEP amplitude to the average change in the electrocardiogram (ECG) amplitude, both across the first significant HEP time window for posterior insula (PI) relative to Sham. X-axis is the change (PI-Sham) in the ECG amplitude (μV) and y-axis is the change (PI-Sham) in the HEP amplitude (μV). Solid red line is the fitted regression line and the dotted lines represent the 95% confidence bounds. P = p-value. (Right) Linear regression comparing the average change in HEP and ECG amplitudes across the second significant HEP time window for PI relative to dorsal anterior cingulate (dACC) conditions. **C.** (Left) Linear regression comparing the average change in the HEP amplitude across the first significant HEP time window to the average change in amplitude across the significant cardiac field artifact (CFA) window for PI relative to Sham conditions. X-axis is the change (PI-Sham) in the CFA amplitude (μV) and y-axis is the change (PI-Sham) in the HEP amplitude (μV). Solid red line is the fitted regression line and the dotted lines represent the 95% confidence interval. P = p-value. (Right) Linear regression comparing the average change in HEP amplitudes across the second significant HEP time window and to the average change in amplitude across the significant CFA time period for PI relative to dACC conditions.

As an additional check to ensure changes in the ECG amplitudes were not driving observed changes in the HEP, we employed linear regression to compare the observed differences in HEP to differences in ECG amplitudes averaged across each significant time window. For the first significant window within the TOI of the HEP, changes in ECG amplitudes (PI-Sham) did not predict changes in the HEP amplitudes (PI-Sham) (R^2^_adj_ = 0.06, F(1,14) = 2.03, p = 0.18) (**Figure 4B left**). For the second significant window within the TOI of the HEP, ECG amplitudes (PI-dACC) did not predict changes in the HEP amplitudes (PI-dACC) (R^2^_adj_ = -0.06, F(1,14) = 0.16, p = 0.69) (**Figure 4B right**).

In the permutation analysis (**Figure 3A**), there were four consecutive milliseconds within the CFA showing significant group differences from 33 to 36ms post R-peak (all p-values < 0.05). This time window was used for control analyses to compare against observed changes in the HEP to check if changes in the significant CFA window predicted changes in either significant window of the HEP. We used the PI-Sham and PI-dACC differences in the HEP and CFA amplitudes averaged across the first and second significant time window, respectively. For the first significant window within the TOI of the HEP, changes in CFA amplitudes (PI-Sham) did not predict changes in the HEP amplitudes (PI-Sham) (R^2^_adj_ = -0.03, F(1,14) = 0.52, p = 0.48) (**Figure 4C left**). For the second significant window within the TOI of the HEP, changes in CFA amplitudes (PI-dACC) did not predict changes in the HEP amplitudes (PI-dACC) (R^2^_adj_ = -0.01, F(1,14) = 0.82, p = 0.38) (**Figure 4C right**).

### Modelling Effects of Pressure

The mean ± SEM estimated intracranial LIFU pressure across conditions was 189.72 ± 21.21, 276.58 ± 23.04, and 115.05 ± 13.25 for AI, PI, and dACC conditions, respectively. A one-way ANOVA demonstrated a significant difference between conditions (F = 20.42 (2,30), p-value < 0.001). Post-hoc testing revealed the PI was significantly higher than both AI and dACC while the AI was significantly higher than the dACC (**Figure 5A**). To examine how pressure and condition explain observed differences in the HEP, we employed linear mixed effects modelling to predict changes in HEP amplitudes with models including either condition, pressure, condition and pressure, or condition, pressure, and the condition-by-pressure interactions as fixed effects. For the first time window (PI-Sham), the full model (including pressure, condition, and the pressure-by-condition interaction) performed significantly better than condition alone (Log likelihood: -91.15 vs. -99.50, LRstat = 16.69(2), p < 0.001) and condition plus pressure with no interaction term (Log likelihood: -91.15 vs. -97.47, LRstat = 6.55(1), p = 0.010). The full model trended towards performing better than pressure alone (Log likelihood: -91.15 vs. -94.02, LRstat = 5.74(2), p = 0.057) and the AIC was smaller for the full model (194.3 vs 196.0), so it was taken as the final model.

**Figure 5.**
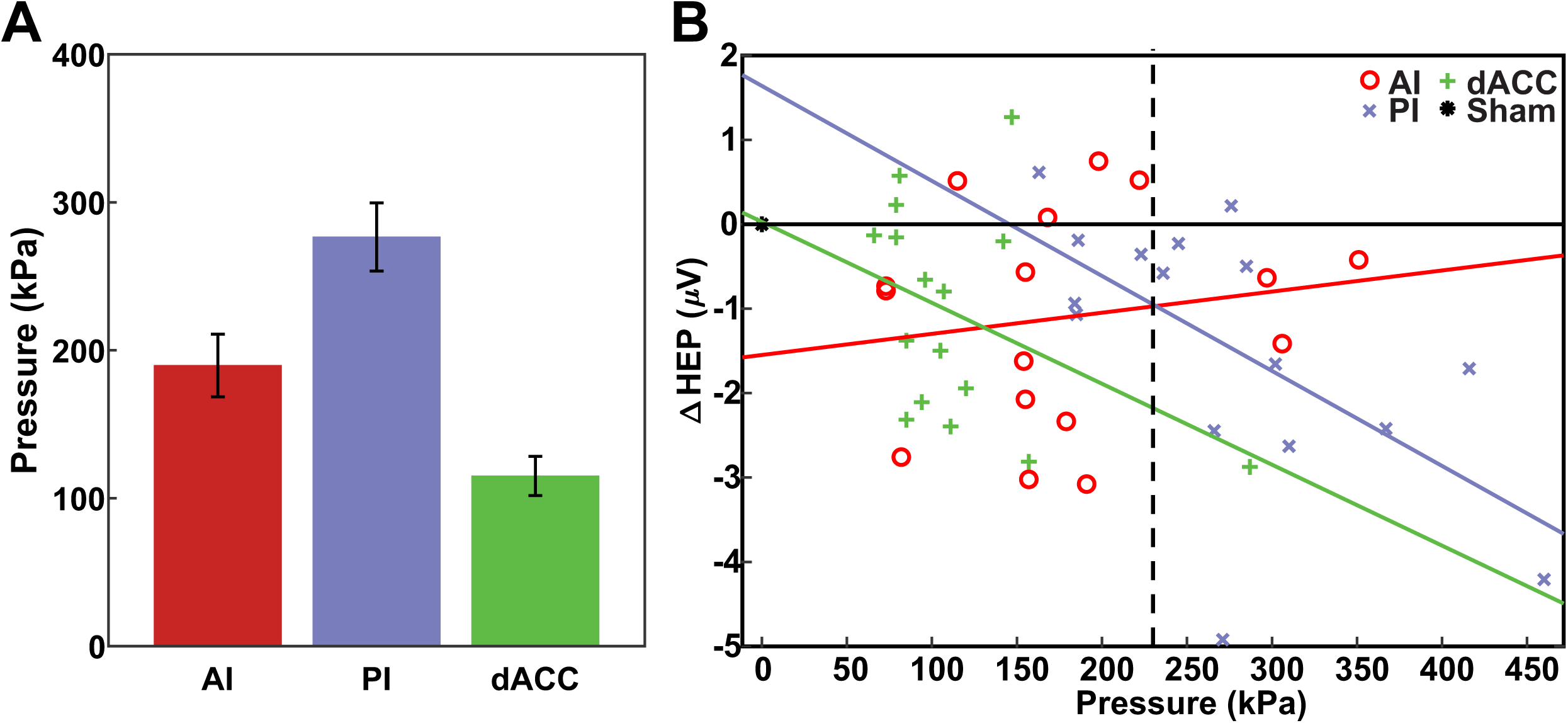
Modelling the effects of pressure on observed HEP differences. **A.** Group (N=16) mean ± SEM estimated intracranial LIFU pressure in kPa across conditions. **B.** Visualization of the significant condition-by-pressure interaction identified during linear mixed effects modelling for the first significant heartbeat-evoked potential (HEP) time window. X-axis is pressure in kilopascals (kPa) and the y-axis is the change in the HEP amplitudes in microvolts (μV). AI = anterior insula (red); PI = posterior insula (purple); dACC = dorsal anterior cingulate cortex (green).

The full model significantly predicted HEP amplitudes (F(3,60) = 23.73, p < 0.001, R^2^_adj_ = 0.52) (**Table 1**). Analysis of individual covariates revealed a significant effect of the condition-by-pressure interaction (t(60) = -2.45, p = 0.017) (**Figures 5B & Table 1**). Both LIFU to the PI and dACC demonstrated an inverse relationship between pressure and HEP amplitude, i.e. higher pressure predicts more negative HEP amplitudes relative to Sham (**Figure 5B**).

**Table 1.**
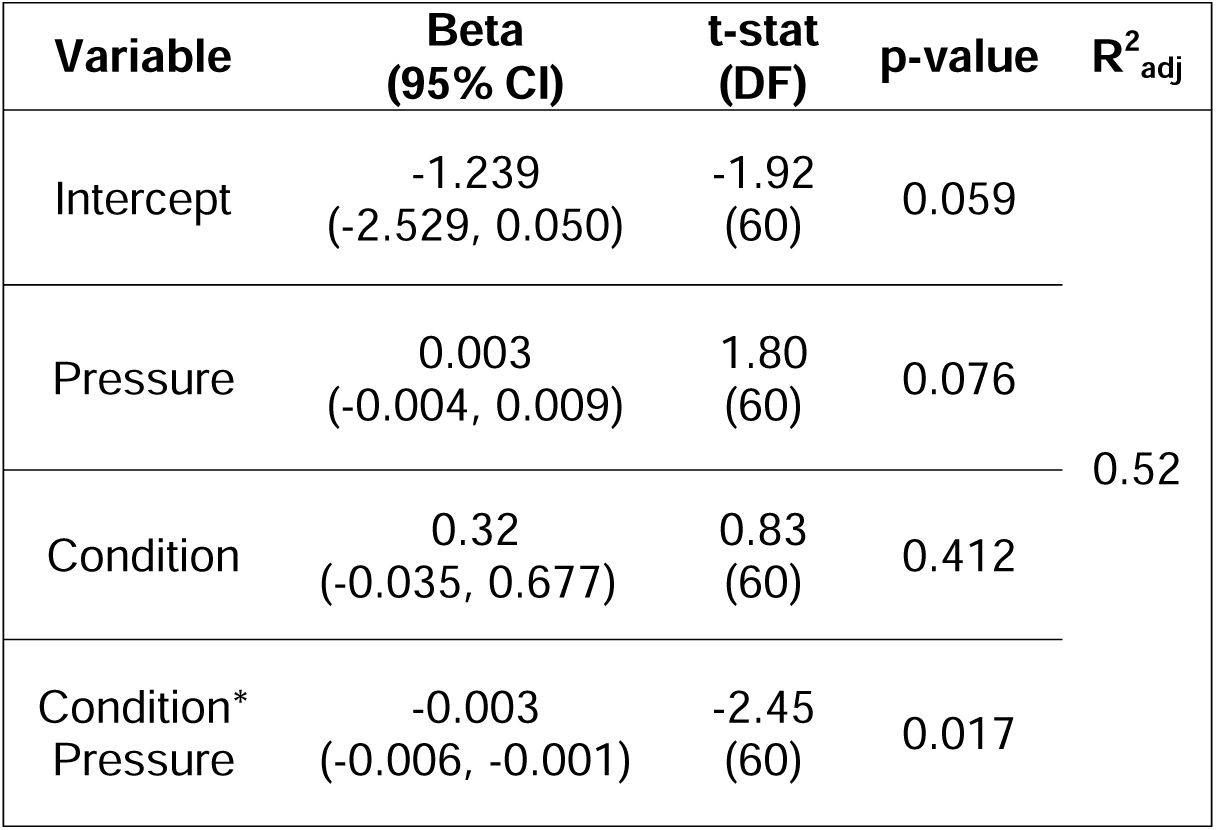
Linear mixed effects model results. Results of the linear mixed effects model which included variables for the main effects of condition and pressure as well as the condition-by-pressure interaction. CI = confidence interval; DF = degrees of freedom; R^2^_adj_ = adjusted r-squared.

For the second time window (PI-dACC), no model was significantly better according to the theoretical likelihood ratio test and none significantly predicted HEP amplitudes, so was not explored further.

### Heart Rate and Heart Rate Variability

The median (IQR) heart rate during LIFU application was 68.59 (16.8), 68.17 (14.64), 71.34 (12.85), and 68.64 (13.62) for AI, PI, dACC, and Sham conditions, respectively. A kruskal-wallis test revealed no significant differences between conditions (chi-square = 0.94 (3,60), p-value = 0.81, η^2^ = 0.01) (**Figure 6A**).

**Figure 6.**
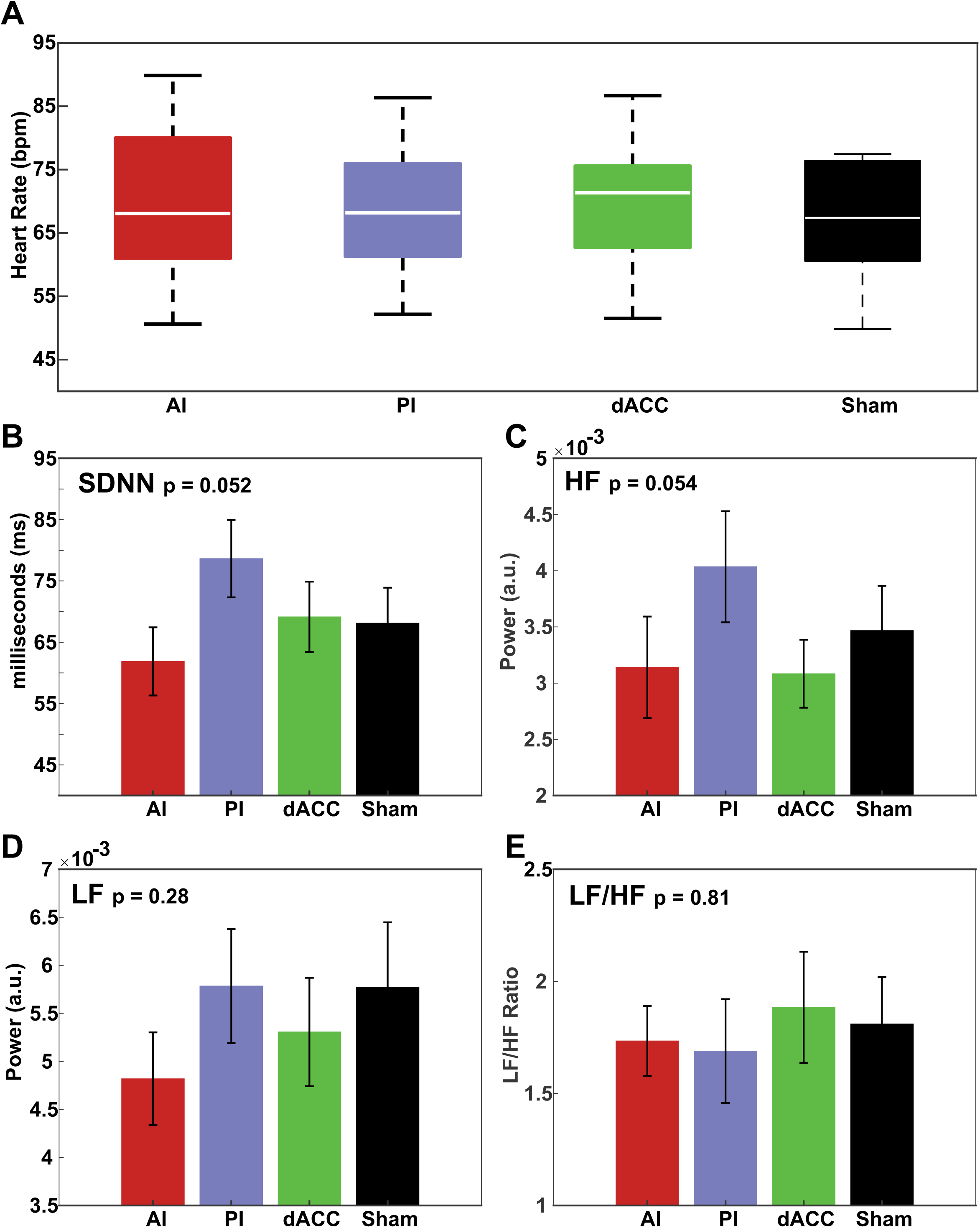
Effects of LIFU on Heart Rate & Heart-rate variability. **A.** Group (N=16) median and interquartile range for heart rate across conditions. Y-axis is heart rate in beats per minute (bpm). **B.** Group (N=16) mean ± SEM standard deviation of normal sinus beats (SDNN) across conditions. Y-axis is time in milliseconds (ms). **C.** Group (N=16) mean ± SEM high frequency (HF) power across conditions. Y-axis is power in absolute units (a.u.). **D.** Group (N=16) mean ± SEM low frequency (LF) power across conditions. Y-axis is power in a.u. **E.** Group (N=16) mean ± SEM LF/HF ratio across conditions. None met statistical significance.

The mean ± SEM SDNN (ms) during LIFU was 61.88 ± 5.56, 78.64 ± 6.31, 69.15 ± 5.73, and 68.11 ± 5.78 for AI, PI, dACC, and Sham conditions, respectively. A one-way ANOVA revealed a trend towards significant differences between conditions (F(3,45) = 2.78, p = 0.052, η^2^ = 0.07) (**Figure 6B**).

The mean ± SEM HF power (absolute units or a.u.) during LIFU was 0.0031 ± 0.0005, 0.0040 ± 0.0005, 0.0031 ± 0.0003, and 0.0035 ± 0.0004 for AI, PI, dACC, and Sham conditions, respectively. A one-way ANOVA revealed a trend towards significant differences between conditions (F(3,45) = 2.75, p = 0.054, η^2^ = 0.05) (**Figure 6C**).

The mean ± SEM LF power (a.u.) during LIFU was 0.0048 ± 0.0005, 0.0058 ± 0.0006, 0.0053 ± 0.0006, and 0.0058 ± 0.0007 for AI, PI, dACC, and Sham conditions, respectively. A one-way ANOVA revealed no significant differences between conditions (F(3,45) = 1.31, p = 0.28, η^2^ = 0.03) (**Figure 6D**).

The mean ± SEM LF/HF ratio during LIFU was 1.73 ± 0.16, 1.69 ± 0.23, 1.88 ± 0.25, and 1.81 ± 0.21 for AI, PI, dACC, and Sham conditions, respectively. A one-way ANOVA revealed no significant differences between conditions (F(3,45) = 0.32, p = 0.81, η^2^ = 0.01) (**Figure 6E**).

### Auditory Masking

For the question “I could hear the LIFU stimulation,” the mean ± SEM score was 1.44 ± 0.48, 2.00 ± 0.55, 2.94 ± 0.56, and 1.94 ± 0.50 for AI, PI, dACC, and Sham conditions, respectively. A kruskal-wallis test revealed no significant difference between conditions (chi-square = 3.85 (3,60), p = 0.28) (**Figure A.1A**).

For the question “I could feel the LIFU stimulation,” the mean ± SEM score was 0.88 ± 0.30, 0.81 ± 0.28, 0.69 ± 0.28, and 0.56 ± 0.22 for AI, PI, dACC, and Sham conditions, respectively. A kruskal-wallis test revealed no significant difference between conditions (chi-square = 0.73 (3,60), p = 0.87) (**Figure A.1B**).

For the question “I believe I experienced LIFU stimulation,” the mean ± SEM score was 3.31 ± 0.41, 3.13 ± 0.33, 2.94 ± 0.40, and 2.13 ± 0.33 for AI, PI, dACC, and Sham conditions, respectively. A kruskal-wallis test revealed no significant difference between conditions (chi-square = 6.7 (3,60), p = 0.08). Since the results for this question were trending, posthoc analysis revealed the effect was driven by differences between AI and Sham which was not the active site that drove any of the above results (**Figure A.1C**).

### Report of Symptoms

The raw symptoms reported before and after LIFU for each condition are presented in **Figure A.2A & B**. The most common symptom before or after any intervention was sleepiness. Prior to LIFU, one subject for both the PI and dACC conditions reported severe sleepiness. This was not present after LIFU application. In addition, we adjusted the symptoms present after LIFU with the symptoms present before LIFU to get an indication of new symptoms following the intervention (**Figure A.2C**). After adjustment, there were no new moderate or severe symptoms across any arm for all queried symptoms. The most common new symptom was sleepiness with 1, 3, 3, and 5 new mild reports for the AI, PI, dACC, and Sham conditions, respectively. A full breakdown of the counts for each symptom can be seen in **Figure A.2**.

## DISCUSSION

In this proof-of-concept study, we investigated the effect of 500 kHz LIFU delivered to either the AI, PI, and dACC compared to Sham on the amplitude of the heartbeat evoked potential, heart rate and heart-rate variability. We identified two distinct time windows in the EEG HEP data with significant group differences that could not be explained by changes in the electrocardiogram or by changes in the cardiac field artifact. These included an early time window around 280ms post R-peak where LIFU to PI was lower than Sham and a later time window around 650ms post R-peak where LIFU to PI was lower than LIFU to dACC. LIFU to the AI and dACC had no effect on the HEP at any time point. We also found no effect of LIFU to any of the targets on mean heart rate or time and frequency domain metrics of heart-rate variability.

### LIFU to the PI attenuates afferent cardiovascular information

We identified a distinct time window in the EEG data around 280ms after the R-peak where LIFU to the PI decreased the HEP amplitude relative to Sham. This change was not related to changes in the CFA or ECG data. Linear mixed modelling showed a condition-by-pressure interaction where increased estimated LIFU pressure at the intracranial target predicted greater decreases in the HEP amplitude for the PI condition. The HEP is commonly considered to reflect cortical processing of afferent cardiac or baroreceptor activity [20,21]. Afferent cardiovascular information the heart and baroreceptors travels to the nucleus of the solitary tract (NTS) where it integrates information with signals from the Lamina I spinothalamic system in the ventrolateral medulla, parabrachial nucleus, periaqueductal gray, and thalamus, before relaying to the insula and dACC [1,2,6]. The insula, often referred to as the primary interoceptive cortex [1,70], receives this Lamina I and NTS information along its posterior and dorsal regions, making the PI the primary site receiving direct interoceptive input [7,12,14,71,72]. In addition, bipolar recordings in the PI during an intracranial study of the HEP in humans have provided evidence the PI is a source of the HEP [29]. The current demonstration of site-specific (i.e., PI) modulation of the HEP within similar time windows identified in previous HEP studies provides additional evidence that the PI does indeed contribute to processing or generation of the observed HEP using surface EEG [21,23,24,29,73].

The HEP looks to be malleable and is modulated by a range of attentional or arousal-based interoceptive tasks [21], including pain [73]. For example, both experimentally-induced pain and chronic pain leads to a concomitant reduction in HEP amplitude that correlated with pain ratings or measures of interoceptive awareness [24,73]. One potential explanation of these results is the presence of pain, whether acute or chronic, overrides or attenuates interoceptive information due to the high saliency of pain. It is possible LIFU to the PI engaged a similar mechanism by directly inhibiting ascending interoceptive information. This would match prior literature that shows the exact same LIFU parameters are inhibitory across a range of brain targets [33,40,41,67].

### LIFU to the AI or dACC did not affect the HEP

We found no evidence for LIFU to the AI or dACC to affect the amplitude of the HEP at any time point. The dACC, similar to the PI, receives direct afferent cardiovascular information from the thalamus but through a distinct nucleus (MDvc) from what supplies the PI (VMpo) [7,12,74,75]. The AI, while not receiving direct afferent thalamic projections, directly communicates with the PI and dACC and serves as an integration hub which helps switch between large-scale brain networks and contributes to efferent visceromotor control [3,14,15,72,76]. Prior intracranial work has demonstrated the HEP can be recorded from the dACC and anterior insula [29]. One potential explanation for the null results is that subregions of the AI and dACC have distinct structural and functional connectivity profiles, thus likely sub-serving different functions [72,77–80]. For example, the ventral AI appears to have a more direct role in efferent visceromotor control compared the dorsal AI region we targeted [14,72]. Other insular targets such as the mid-insula, often grouped with the AI, also appears to play a distinct role in interoceptive processing [81,82]. For the dACC, our target was posterior to sites identified both through intracranial and fMRI studies of the HEP [20,29]. Future studies should aim to precisely map insular and cingulate subregions for effects on the HEP.

### LIFU pressure predicts the response to LIFU in humans

Linear mixed effects modelling demonstrated a role of pressure in the observed effects on the HEP whereby higher pressures produced greater changes in HEP amplitudes. Prior in-vitro and animal work has consistently shown dose-response relationships with LIFU where more energy produces stronger effects on the outcome of interest [83–86]. While previous human studies have reported estimated intracranial energy [41,87–89], the present investigation along with its companion behavioral study [42] provide early evidence of a dose-response to LIFU in humans on measures ranging from pain perception to neurophysiologic responses. Interestingly, in the present study the observed effects of pressure on HEP amplitudes were the same for both PI and dACC conditions, but not for AI. This finding matches our hypotheses that the PI and dACC would have similar effects on HEP amplitudes due to both receiving direct afferent interoceptive information as opposed to the AI [1,7,9,12].

Dose-responses are a valuable measure for assessing causality in any human brain mapping study, and establishing a dose-response relationship for a desired effect at a specific neural target is critical for clinical applications of any non-invasive brain stimulation technique [90]. The differential effect of pressure at each site in the present study helps build this evidence base for LIFU, although more work must be done to establish a minimal effective dose.

### Effects of LIFU on HRV may be state-dependent

We observed trends for changes in the SDNN and HF power during LIFU application, but none reached statistical significance. Previous work using the same LIFU parameters in a similar concurrent design clearly demonstrated effects of LIFU to the insula and dACC on these metrics, yet these studies had LIFU delivered concurrently with a painful stimulus [40,41]. Considering a growing literature points towards state-dependency on the effects of neuromodulation with LIFU [35,91], as well as other brain stimulation techniques [92,93], there may be an interaction between physiologic states and effects of LIFU on HRV. Evaluation of this possibility would benefit from additional studies involving larger sample sizes.

#### Limitations and future directions

There are several limitations to this study. Although this proof-of-concept study demonstrated modulation of the HEP from LIFU to PI, the reliance on a single electrode represents a limitation. Future work is needed to replicate the present findings across a larger number of scalp leads and in a larger sample size. Such studies could utilize data-driven cluster-based permutation approaches [68] to identify electrode clusters showing maximal responses, which could in term support the application of source localization algorithms to analyze the underlying HEP generators. Additionally, our dose-response relationship was based on differences in estimated pressure as opposed to a pre-defined dosing strategy. Establishing clear dose-response relationships require studies explicitly exploring dose-response thus our findings of the effect of LIFU pressure should be considered preliminary. Finally, our choice or targets within the AI, PI, and dACC may not necessarily reflect the optimal targets for investigating the role of these regions in the HEP. Therefore, the null findings in the AI and dACC should not be taken as conclusive evidence of their role in the HEP. More comprehensive mapping of insular and cingulate subregions as well as more advanced brain stimulation targeting approaches such as individualized resting state functional connectivity were not explored and could be used in future studies to better answer questions about these regions’ contribution to the HEP.

## CONCLUSIONS AND FUTURE DIRECTIONS

As a putative biomarker of brain-heart interaction, the HEP may be partially responsive to modulation of the posterior insula by LIFU. Future work should investigate LIFU effects on the HEP using additional electrode leads, at a range of pre-defined doses, and in patient populations to evaluate clinical relevancy.

## Acknowledgements

We would like to thank Jessica Florig for her assistance with data collection and data management.

## APPENDIX A

**Figure A.1.**
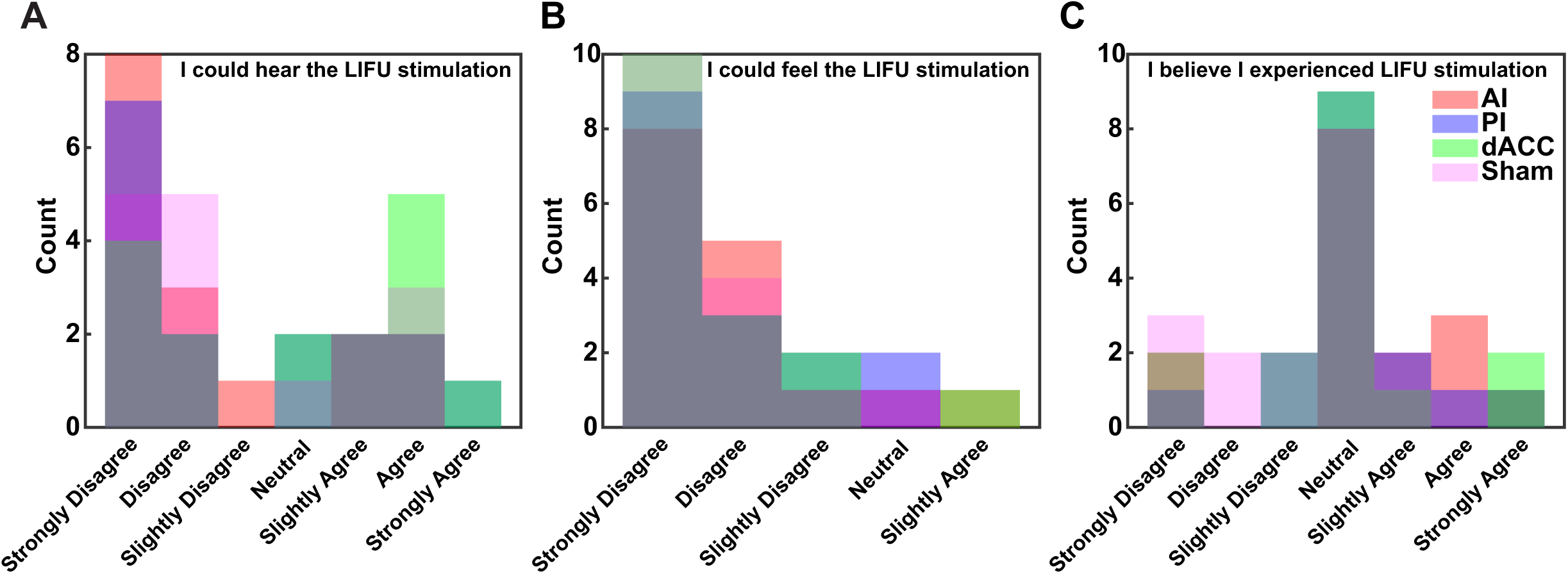
Auditory masking results. Counts from each participant and condition across the 7-point Likert Scale for each question in the auditory masking questionnaire (AMQ). **A.** Results from the question “I could hear the LIFU stimulation.” **B.** Results from the question “I could feel the LIFU stimulation.” **C.** Results from the question “I believe I experienced LIFU stimulation.”

**Figure A.2.**
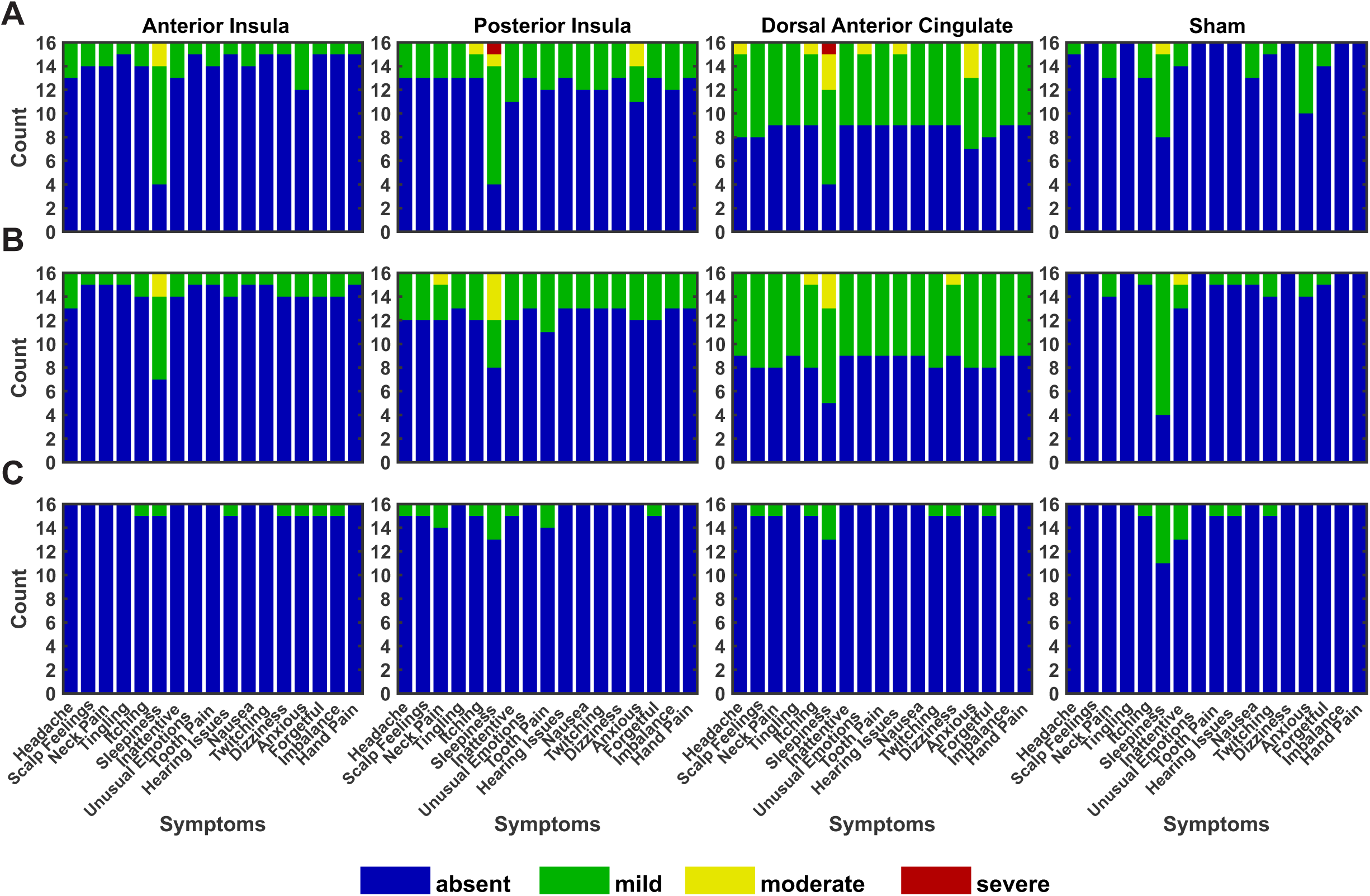
Report of symptoms questionnaire results. Counts for each queried symptom and severity level for each condition. From left to right, the columns represent the anterior insula, posterior insula, dorsal anterior cingulate, and Sham conditions. Severity levels of each symptom are represented as absent (blue), mild (green), moderate (yellow), and severe (red). **A.** Presence of symptoms prior to LIFU or Sham application. **B.** Presence of symptoms after LIFU or Sham application. **C.** Presence of symptoms after LIFU or Sham application adjusted for symptoms present prior to LIFU or Sham application.

**Figure A.3.**
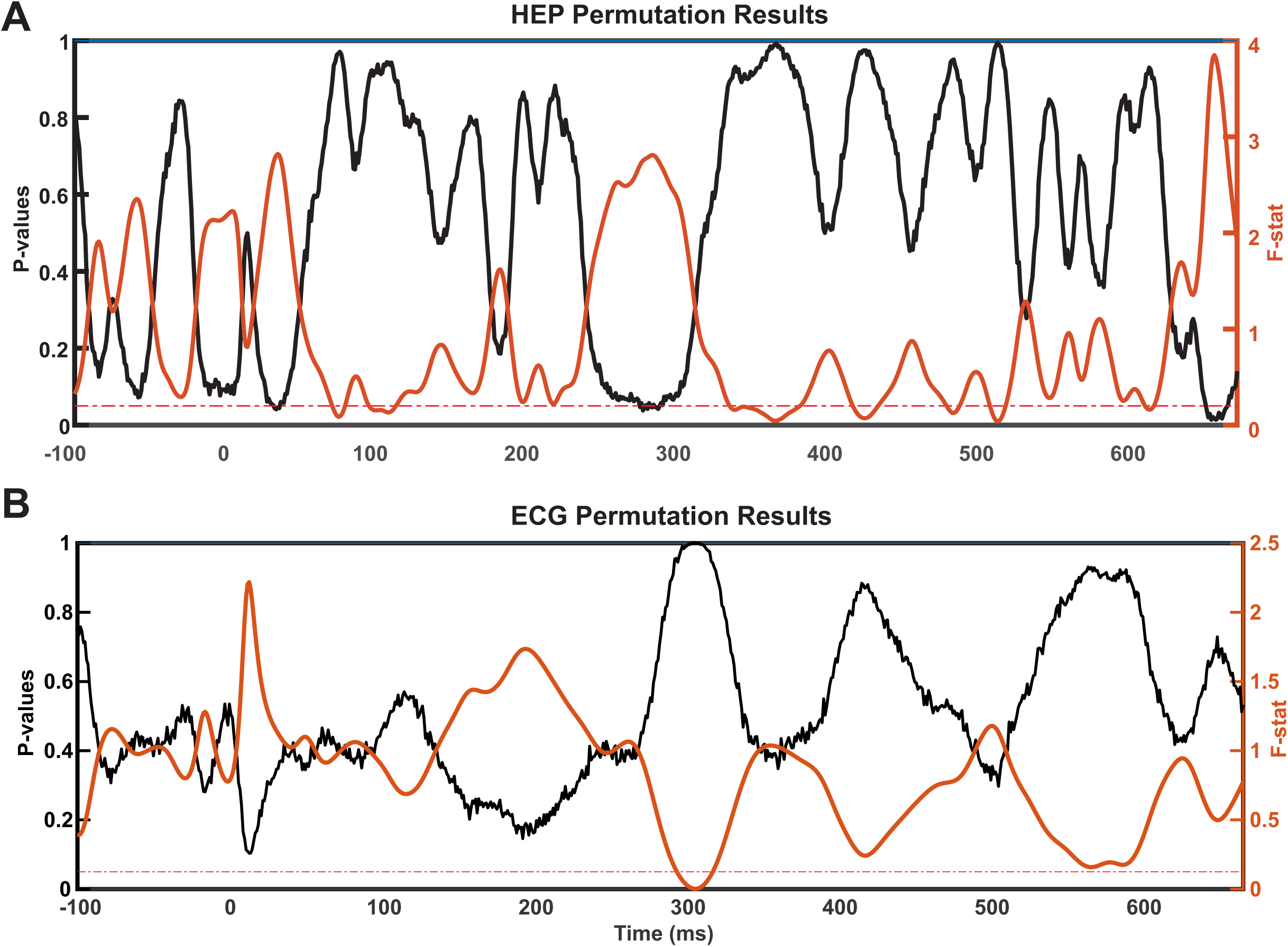
Permutation testing F-statistic and p-value plots. **A.** Results from the permutation analysis across time for the HEP data in Figure 3A. X-axis is time (ms), left y-axis and black line represents p-values, right y-axis and orange line represents F-statistics from one-way ANOVA. Horizontal dashed red line represents p = 0.05. **B.** Results from the permutation analysis across time for ECG data in Figure 4A. X-axis is time (ms), left y-axis and black line represents p-values, right y-axis and orange line represents F-statistics. Horizontal dashed red line represents p = 0.05.

**Table A.1.** Individual target depths. Target depths from the scalp for each target from each individual measured in millimeters (mm).

| Sex | Subject | AI | PI | dACC |
| --- | --- | --- | --- | --- |
| Male | 1 | 40.8 | 42.4 | 49.3 |
|  | 2 | 30.4 | 35.8 | 49.2 |
|  | 3 | 37.7 | 45.0 | 53.7 |
|  | 4 | 30.4 | 35.8 | 54.5 |
|  | 5 | 35.8 | 42.0 | 45.6 |
|  | 6 | 35.8 | 40.7 | 47.8 |
|  | Mean | 35.2 | 40.3 | 50.0 |
|  | SD | 4.1 | 3.7 | 3.4 |
| Female | 1 | 36.3 | 41.9 | 48.8 |
|  | 2 | 34.5 | 38.9 | 46.0 |
|  | 3 | 33.6 | 36.9 | 47.0 |
|  | 4 | 31.2 | 37.3 | 43.8 |
|  | 5 | 33.4 | 38.1 | 46.7 |
|  | 6 | 34.7 | 39.7 | 45.7 |
|  | 7 | 36.2 | 41.2 | 45.7 |
|  | 8 | 34.3 | 38.1 | 42.7 |
|  | 9 | 34.0 | 37.2 | 41.5 |
|  | 10 | 32.3 | 38.8 | 46.6 |
|  | Mean | 34.1 | 38.8 | 45.5 |
|  | SD | 1.6 | 1.7 | 2.2 |
| Total | Mean | 34.5 | 39.4 | 47.3 |
|  | SD | 2.6 | 2.6 | 3.4 |

## REFERENCES

[1] Craig AD (Bud). How do you feel? Interoception: the sense of the physiological condition of the body. Nat Rev Neurosci 2002;3:655–66. 10.1038/nrn894.

[2] Berntson GG, Khalsa SS. Neural Circuits of Interoception. Trends Neurosci 2021;44:17–28. 10.1016/j.tins.2020.09.011.

[3] Khalsa SS, Adolphs R, Cameron OG, Critchley HD, Davenport PW, Feinstein JS, et al. Interoception and Mental Health: A Roadmap. Biological Psychiatry: Cognitive Neuroscience and Neuroimaging 2018;3:501–13. 10.1016/j.bpsc.2017.12.004.

[4] Bonaz B, Lane RD, Oshinsky ML, Kenny PJ, Sinha R, Mayer EA, et al. Diseases, Disorders, and Comorbidities of Interoception. Trends in Neurosciences 2021;44:39–51. 10.1016/j.tins.2020.09.009.

[5] Di Lernia D, Serino S, Riva G. Pain in the body. Altered interoception in chronic pain conditions: A systematic review. Neuroscience & Biobehavioral Reviews 2016;71:328–41. 10.1016/j.neubiorev.2016.09.015.

[6] Craig AD (Bud). How do you feel — now? The anterior insula and human awareness. Nat Rev Neurosci 2009;10:59–70. 10.1038/nrn2555.

[7] Dum RP, Levinthal DJ, Strick PL. The Spinothalamic System Targets Motor and Sensory Areas in the Cerebral Cortex of Monkeys. J Neurosci 2009;29:14223–35. 10.1523/JNEUROSCI.3398-09.2009.

[8] Gasparini S, Howland JM, Thatcher AJ, Geerling JC. Central afferents to the nucleus of the solitary tract in rats and mice. Journal of Comparative Neurology 2020;528:2708–28. 10.1002/cne.24927.

[9] Palma J-A, Benarroch EE. Neural control of the heart: Recent concepts and clinical correlations. Neurology 2014;83:261–71. 10.1212/WNL.0000000000000605.

[10] Mandonnet V, Obaid S, Descoteaux M, St-Onge E, Devaux B, Levé C, et al. Electrostimulation of the white matter of the posterior insula and medial operculum: perception of vibrations, heat, and pain. Pain 2024;165:565–72. 10.1097/j.pain.0000000000003069.

[11] Craig ADB. Topographically organized projection to posterior insular cortex from the posterior portion of the ventral medial nucleus in the long-tailed macaque monkey. J Comp Neurol 2014;522:36–63. 10.1002/cne.23425.

[12] Craig AD (Bud). Distribution of trigeminothalamic and spinothalamic lamina I terminations in the macaque monkey. Journal of Comparative Neurology 2004;477:119–48. 10.1002/cne.20240.

[13] Seth AK. Interoceptive inference, emotion, and the embodied self. Trends in Cognitive Sciences 2013;17:565–73. 10.1016/j.tics.2013.09.007.

[14] Barrett LF, Simmons WK. Interoceptive predictions in the brain | Nature Reviews Neuroscience. Nature Reviews Neuroscience 2015;16:419–29. 10.1038/nrn3950.

[15] Seeley WW. The Salience Network: A Neural System for Perceiving and Responding to Homeostatic Demands. J Neurosci 2019;39:9878–82. 10.1523/JNEUROSCI.1138-17.2019.

[16] Beissner F, Meissner K, Bär K-J, Napadow V. The Autonomic Brain: An Activation Likelihood Estimation Meta-Analysis for Central Processing of Autonomic Function. J Neurosci 2013;33:10503–11. 10.1523/JNEUROSCI.1103-13.2013.

[17] Siddiqi SH, Schaper FLWVJ, Horn A, Hsu J, Padmanabhan JL, Brodtmann A, et al. Brain stimulation and brain lesions converge on common causal circuits in neuropsychiatric disease. Nat Hum Behav 2021;5:1707–16. 10.1038/s41562-021-01161-1.

[18] Taylor JJ, Lin C, Talmasov D, Ferguson MA, Schaper FLWVJ, Jiang J, et al. A transdiagnostic network for psychiatric illness derived from atrophy and lesions. Nat Hum Behav 2023:1–10. 10.1038/s41562-022-01501-9.

[19] Wager TD, Atlas LY, Lindquist MA, Roy M, Woo C-W, Kross E. An fMRI-Based Neurologic Signature of Physical Pain. N Engl J Med 2013;368:1388–97. 10.1056/NEJMoa1204471.

[20] Park H-D, Blanke O. Heartbeat-evoked cortical responses: Underlying mechanisms, functional roles, and methodological considerations. NeuroImage 2019;197:502–11. 10.1016/j.neuroimage.2019.04.081.

[21] Coll M-P, Hobson H, Bird G, Murphy J. Systematic review and meta-analysis of the relationship between the heartbeat-evoked potential and interoception. Neuroscience & Biobehavioral Reviews 2021;122:190–200. 10.1016/j.neubiorev.2020.12.012.

[22] Schandry R, Sparrer B, Weitkunat R. From the heart to the brain: A study of heartbeat contingent scalp potentials. International Journal of Neuroscience 1986;30:261–75. 10.3109/00207458608985677.

[23] Engelen T, Solcà M, Tallon-Baudry C. Interoceptive rhythms in the brain. Nat Neurosci 2023:1–15. 10.1038/s41593-023-01425-1.

[24] Solcà M, Park H-D, Bernasconi F, Blanke O. Behavioral and neurophysiological evidence for altered interoceptive bodily processing in chronic pain. NeuroImage 2020;217:116902. 10.1016/j.neuroimage.2020.116902.

[25] Terhaar J, Viola FC, Bär K-J, Debener S. Heartbeat evoked potentials mirror altered body perception in depressed patients. Clinical Neurophysiology 2012;123:1950–7. 10.1016/j.clinph.2012.02.086.

[26] Pang J, Tang X, Li H, Hu Q, Cui H, Zhang L, et al. Altered Interoceptive Processing in Generalized Anxiety Disorder—A Heartbeat-Evoked Potential Research. Frontiers in Psychiatry 2019;10.

[27] Verdonk C, Teed AR, White EJ, Ren X, Stewart JL, Paulus MP, et al. Heartbeat-evoked neural response abnormalities in generalized anxiety disorder during peripheral adrenergic stimulation. Neuropsychopharmacol 2024:1–9. 10.1038/s41386-024-01806-5.

[28] Quadt L, Critchley HD, Garfinkel SN. The neurobiology of interoception in health and disease. Annals of the New York Academy of Sciences 2018;1428:112–28. 10.1111/nyas.13915.

[29] Park H-D, Bernasconi F, Salomon R, Tallon-Baudry C, Spinelli L, Seeck M, et al. Neural Sources and Underlying Mechanisms of Neural Responses to Heartbeats, and their Role in Bodily Self-consciousness: An Intracranial EEG Study. Cerebral Cortex 2018;28:2351–64. 10.1093/cercor/bhx136.

[30] Mazzola L, Mauguière F, Chouchou F. Central control of cardiac activity as assessed by intra-cerebral recordings and stimulations. Neurophysiologie Clinique 2023;53:102849. 10.1016/j.neucli.2023.102849.

[31] Tyler WJ, Lani SW, Hwang GM. Ultrasonic modulation of neural circuit activity. Current Opinion in Neurobiology 2018;50:222–31. 10.1016/j.conb.2018.04.011.

[32] Darmani G, Bergmann TO, Butts Pauly K, Caskey CF, de Lecea L, Fomenko A, et al. Non-invasive transcranial ultrasound stimulation for neuromodulation. Clin Neurophysiol 2022;135:51–73. 10.1016/j.clinph.2021.12.010.

[33] Legon W, Sato TF, Opitz A, Mueller J, Barbour A, Williams A, et al. Transcranial focused ultrasound modulates the activity of primary somatosensory cortex in humans. Nat Neurosci 2014;17:322–9. 10.1038/nn.3620.

[34] Legon W, Ai L, Bansal P, Mueller JK. Neuromodulation with single-element transcranial focused ultrasound in human thalamus. Hum Brain Mapp 2018;39:1995–2006. 10.1002/hbm.23981.

[35] Yaakub SN, White TA, Roberts J, Martin E, Verhagen L, Stagg CJ, et al. Transcranial focused ultrasound-mediated neurochemical and functional connectivity changes in deep cortical regions in humans | Nature Communications. Nature Communications 2023;14:5318. 10.1038/s41467-023-40998-0.

[36] Butler CR, Rhodes E, Blackmore J, Cheng X, Peach RL, Veldsman M, et al. Transcranial ultrasound stimulation to human middle temporal complex improves visual motion detection and modulates electrophysiological responses. Brain Stimulation: Basic, Translational, and Clinical Research in Neuromodulation 2022;15:1236–45. 10.1016/j.brs.2022.08.022.

[37] Nakajima K, Osada T, Ogawa A, Tanaka M, Oka S, Kamagata K, et al. A causal role of anterior prefrontal-putamen circuit for response inhibition revealed by transcranial ultrasound stimulation in humans. Cell Reports 2022;40:111197. 10.1016/j.celrep.2022.111197.

[38] Mueller J, Legon W, Opitz A, Sato TF, Tyler WJ. Transcranial Focused Ultrasound Modulates Intrinsic and Evoked EEG Dynamics. Brain Stimulation 2014;7:900–8. 10.1016/j.brs.2014.08.008.

[39] Badran BW, Caulfield KA, Stomberg-Firestein S, Summers PM, Dowdle LT, Savoca M, et al. Sonication of the anterior thalamus with MRI-Guided transcranial focused ultrasound (tFUS) alters pain thresholds in healthy adults: A double-blind, sham-controlled study. Brain Stimulation 2020;13:1805–12. 10.1016/j.brs.2020.10.007.

[40] Legon W, Strohman A, In A, Payne B. Noninvasive neuromodulation of subregions of the human insula differentially affect pain processing and heart-rate variability: a within-subjects pseudo-randomized trial. PAIN 2024:10.1097/j.pain.0000000000003171. 10.1097/j.pain.0000000000003171.

[41] Strohman A, Payne B, In A, Stebbins K, Legon W. Low-intensity focused ultrasound to the human dorsal anterior cingulate attenuates acute pain perception and autonomic responses. J Neurosci 2024. 10.1523/JNEUROSCI.1011-23.2023.

[42] In A, Strohman A, Payne B, Legon W. Low-intensity focused ultrasound to the insula and dorsal anterior cingulate has site-specific and pressure dependent effects on pain during measures of central sensitization 2024:2024.01.10.575098. 10.1101/2024.01.10.575098.

[43] Kemp AH, Brunoni AR, Santos IS, Nunes MA, Dantas EM, Carvalho de Figueiredo R, et al. Effects of Depression, Anxiety, Comorbidity, and Antidepressants on Resting-State Heart Rate and Its Variability: An ELSA-Brasil Cohort Baseline Study. AJP 2014;171:1328–34. 10.1176/appi.ajp.2014.13121605.

[44] Kemp AH, Quintana DS, Gray MA, Felmingham KL, Brown K, Gatt JM. Impact of Depression and Antidepressant Treatment on Heart Rate Variability: A Review and Meta-Analysis. Biological Psychiatry 2010;67:1067–74. 10.1016/j.biopsych.2009.12.012.

[45] Zebhauser PT, Hohn VD, Ploner M. Resting-state electroencephalography and magnetoencephalography as biomarkers of chronic pain: a systematic review. Pain 2023;164:1200–21. 10.1097/j.pain.0000000000002825.

[46] Samuel N, Zeng K, Harmsen IE, Ding MYR, Darmani G, Sarica C, et al. Multi-modal investigation of transcranial ultrasound-induced neuroplasticity of the human motor cortex. Brain Stimulation 2022;15:1337–47. 10.1016/j.brs.2022.10.001.

[47] Hobday DI, Hobson AR, Sarkar S, Furlong PL, Thompson DG, Aziz Q. Cortical processing of human gut sensation: an evoked potential study. Am J Physiol Gastrointest Liver Physiol 2002;283:G335–339. 10.1152/ajpgi.00230.2001.

[48] Aziz Q, Furlong PL, Barlow J, Hobson A, Alani S, Bancewicz J, et al. Topographic mapping of cortical potentials evoked by distension of the human proximal and distal oesophagus. Electroencephalogr Clin Neurophysiol 1995;96:219–28. 10.1016/0168-5597(94)00297-r.

[49] Hobson AR, Khan RW, Sarkar S, Furlong PL, Aziz Q. Development of esophageal hypersensitivity following experimental duodenal acidification. Am J Gastroenterol 2004;99:813–20. 10.1111/j.1572-0241.2004.04167.x.

[50] Strohman A, In A, Stebbins K, Legon W. Evaluation of a Novel Acoustic Coupling Medium for Human Low-Intensity Focused Ultrasound Neuromodulation Applications. Ultrasound in Medicine & Biology 2023. 10.1016/j.ultrasmedbio.2023.02.003.

[51] Yarkoni T, Poldrack RA, Nichols TE, Van Essen DC, Wager TD. Large-scale automated synthesis of human functional neuroimaging data. Nat Methods 2011;8:665–70. 10.1038/nmeth.1635.

[52] Wager TD, Atlas LY, Lindquist MA, Roy M, Woo C-W, Kross E. An fMRI-Based Neurologic Signature of Physical Pain. N Engl J Med 2013;368:1388–97. 10.1056/NEJMoa1204471.

[53] Liang W, Guo H, Mittelstein DR, Shapiro MG, Shimojo S, Shehata MH. Auditory Mondrian masks the airborne-auditory artifact of focused ultrasound stimulation in humans. Brain Stimulation: Basic, Translational, and Clinical Research in Neuromodulation 2023;0. 10.1016/j.brs.2023.03.002.

[54] Legon W, Adams S, Bansal P, Patel PD, Hobbs L, Ai L, et al. A retrospective qualitative report of symptoms and safety from transcranial focused ultrasound for neuromodulation in humans. Scientific Reports 2020;10:1–10.

[55] Legon W, Strohman A, In A, Stebbins K, Payne B. Non-invasive neuromodulation of sub-regions of the human insula differentially affect pain processing and heart-rate variability 2023:2023.05.05.539593. 10.1101/2023.05.05.539593.

[56] Petzschner FH, Weber LA, Wellstein KV, Paolini G, Do CT, Stephan KE. Focus of attention modulates the heartbeat evoked potential. NeuroImage 2019;186:595–606. 10.1016/j.neuroimage.2018.11.037.

[57] Widmann A, Schröger E, Maess B. Digital filter design for electrophysiological data – a practical approach. Journal of Neuroscience Methods 2015;250:34–46. 10.1016/j.jneumeth.2014.08.002.

[58] Tanner D, Morgan-Short K, Luck SJ. How inappropriate high-pass filters can produce artifactual effects and incorrect conclusions in ERP studies of language and cognition. Psychophysiology 2015;52:997–1009. 10.1111/psyp.12437.

[59] Gray MA, Minati L, Paoletti G, Critchley HD. Baroreceptor activation attenuates attentional effects on pain-evoked potentials. Pain 2010;151:853–61. 10.1016/j.pain.2010.09.028.

[60] Kern M, Aertsen A, Schulze-Bonhage A, Ball T. Heart cycle-related effects on event-related potentials, spectral power changes, and connectivity patterns in the human ECoG. Neuroimage 2013;81:178–90. 10.1016/j.neuroimage.2013.05.042.

[61] Treeby BE, Cox BT. k-Wave: MATLAB toolbox for the simulation and reconstruction of photoacoustic wave fields. JBO 2010;15:021314. 10.1117/1.3360308.

[62] Legon W, Ai L, Bansal P, Mueller JK. Neuromodulation with single-element transcranial focused ultrasound in human thalamus. Human Brain Mapping 2018;39:1995–2006.

[63] Mueller JK, Ai L, Bansal P, Legon W. Computational exploration of wave propagation and heating from transcranial focused ultrasound for neuromodulation. Journal of Neural Engineering 2016;13:056002.

[64] Aubry J-F, Tanter M, Pernot M, Thomas J-L, Fink M. Experimental demonstration of noninvasive transskull adaptive focusing based on prior computed tomography scans. The Journal of the Acoustical Society of America 2003;113:84–93. 10.1121/1.1529663.

[65] Marquet F, Boch A-L, Pernot M, Montaldo G, Seilhean D, Fink M, et al. Non-invasive ultrasonic surgery of the brain in non-human primates. The Journal of the Acoustical Society of America 2013;134:1632–9. 10.1121/1.4812888.

[66] Mueller JK, Ai L, Bansal P, Legon W. Numerical evaluation of the skull for human neuromodulation with transcranial focused ultrasound. J Neural Eng 2017;14:066012. 10.1088/1741-2552/aa843e.

[67] Legon W, Bansal P, Tyshynsky R, Ai L, Mueller JK. Transcranial focused ultrasound neuromodulation of the human primary motor cortex. Scientific Reports 2018;8:1–14.

[68] Maris E, Oostenveld R. Nonparametric statistical testing of EEG- and MEG-data. J Neurosci Methods 2007;164:177–90. 10.1016/j.jneumeth.2007.03.024.

[69] Shaffer F, Ginsberg JP. An Overview of Heart Rate Variability Metrics and Norms. Frontiers in Public Health 2017;5.

[70] Craig AD. A new view of pain as a homeostatic emotion. Trends in Neurosciences 2003;26:303–7. 10.1016/S0166-2236(03)00123-1.

[71] Uddin LQ, Nomi JS, Hébert-Seropian B, Ghaziri J, Boucher O. Structure and Function of the Human Insula: Journal of Clinical Neurophysiology 2017;34:300–6. 10.1097/WNP.0000000000000377.

[72] Kleckner IR, Zhang J, Touroutoglou A, Chanes L, Xia C, Simmons WK, et al. Evidence for a large-scale brain system supporting allostasis and interoception in humans. Nat Hum Behav 2017;1:1– 14. 10.1038/s41562-017-0069.

[73] Shao S, Shen K, Wilder-Smith EPV, Li X. Effect of pain perception on the heartbeat evoked potential. Clinical Neurophysiology 2011;122:1838–45. 10.1016/j.clinph.2011.02.014.

[74] Craig AD. Pain mechanisms: Labeled lines versus convergence in central processing. Annual Review of Neuroscience 2003;26:1–30.

[75] Blomqvist A, Evrard HC. The thalamic projection of pain sensations to the posterior dorsal fundus in the insula: comment on Mandonnet et al. Pain 2024;165:e15–6. 10.1097/j.pain.0000000000003164.

[76] Frot M, Faillenot I, Mauguière F. Processing of nociceptive input from posterior to anterior insula in humans. Human Brain Mapping 2014;35:5486–99. 10.1002/hbm.22565.

[77] Cormie MA, Kaya B, Hadjis GE, Mouseli P, Moayedi M. Insula-cingulate structural and functional connectivity: an ultra-high field MRI study. Cerebral Cortex 2023:bhad244. 10.1093/cercor/bhad244.

[78] Ghaziri J, Tucholka A, Girard G, Houde J-C, Boucher O, Gilbert G, et al. The Corticocortical Structural Connectivity of the Human Insula. Cerebral Cortex 2017;27:1216–28. 10.1093/cercor/bhv308.

[79] Ghaziri J, Tucholka A, Girard G, Boucher O, Houde J-C, Descoteaux M, et al. Subcortical structural connectivity of insular subregions. Sci Rep 2018;8:8596. 10.1038/s41598-018-26995-0.

[80] Beckmann M, Johansen-Berg H, Rushworth MFS. Connectivity-Based Parcellation of Human Cingulate Cortex and Its Relation to Functional Specialization. J Neurosci 2009;29:1175–90. 10.1523/JNEUROSCI.3328-08.2009.

[81] Adamic EM, Teed AR, Avery JA, Cruz F de la, Khalsa SS. Hemispheric divergence of interoceptive processing across psychiatric disorders. eLife 2024;13. 10.7554/eLife.92820.1.

[82] Paulus MP, Khalsa SS. When You Don’t Feel Right Inside: Homeostatic Dysregulation and the Mid-Insular Cortex in Psychiatric Disorders. AJP 2021;178:683–5. 10.1176/appi.ajp.2021.21060622.

[83] Tufail Y, Matyushov A, Baldwin N, Tauchmann ML, Georges J, Yoshihiro A, et al. Transcranial Pulsed Ultrasound Stimulates Intact Brain Circuits. Neuron 2010;66:681–94. 10.1016/j.neuron.2010.05.008.

[84] Yoo S, Mittelstein DR, Hurt RC, Lacroix J, Shapiro MG. Focused ultrasound excites cortical neurons via mechanosensitive calcium accumulation and ion channel amplification. Nat Commun 2022;13:493. 10.1038/s41467-022-28040-1.

[85] Mohammadjavadi M, Ye PP, Xia A, Brown J, Popelka G, Pauly KB. Elimination of peripheral auditory pathway activation does not affect motor responses from ultrasound neuromodulation. Brain Stimulation 2019;12:901–10. 10.1016/j.brs.2019.03.005.

[86] Murphy KR, Farrell JS, Gomez JL, Stedman QG, Li N, Leung SA, et al. A tool for monitoring cell type– specific focused ultrasound neuromodulation and control of chronic epilepsy. Proceedings of the National Academy of Sciences 2022;119:e2206828119. 10.1073/pnas.2206828119.

[87] Legon W, Strohman A, In A, Stebbins K, Payne B. Non-invasive neuromodulation of sub-regions of the human insula differentially affect pain processing and heart-rate variability. bioRxiv 2023:2023.05.05.539593. 10.1101/2023.05.05.539593.

[88] Lee W, Kim H, Jung Y, Song I-U, Chung YA, Yoo S-S. Image-Guided Transcranial Focused Ultrasound Stimulates Human Primary Somatosensory Cortex. Sci Rep 2015;5:8743. 10.1038/srep08743.

[89] Lee W, Kim H-C, Jung Y, Chung YA, Song I-U, Lee J-H, et al. Transcranial focused ultrasound stimulation of human primary visual cortex. Sci Rep 2016;6:34026. 10.1038/srep34026.

[90] Siddiqi SH, Kording KP, Parvizi J, Fox MD. Causal mapping of human brain function. Nat Rev Neurosci 2022;23:361–75. 10.1038/s41583-022-00583-8.

[91] Yang P-F, Phipps MA, Jonathan S, Newton AT, Byun N, Gore JC, et al. Bidirectional and state-dependent modulation of brain activity by transcranial focused ultrasound in non-human primates. Brain Stimulation: Basic, Translational, and Clinical Research in Neuromodulation 2021;14:261–72. 10.1016/j.brs.2021.01.006.

[92] Bradley C, Nydam AS, Dux PE, Mattingley JB. State-dependent effects of neural stimulation on brain function and cognition. Nat Rev Neurosci 2022. 10.1038/s41583-022-00598-1.

[93] Scangos KW, Makhoul GS, Sugrue LP, Chang EF, Krystal AD. State-dependent responses to intracranial brain stimulation in a patient with depression | Nature Medicine. Nature Medicine 2021;27:229–31. 10.1038/s41591-020-01175-8.

